# Chemokine and opioid peptide scavenging through constitutive and ligand-induced release of ACKR3-bearing extracellular vesicles

**DOI:** 10.64898/2026.06.18.733119

**Authors:** Christie B. Palmer, Lucie Rospape, Max Meyrath, Caitrín Crudden, Manuel Counson, Anne-Katrin, Lotte Di Niro, Ana Alonso Bartolome, Cláudio Pinheiro, Vanessa Klapp, Ester Cassano, Stéphane A. Laporte, Stephen J. Hill, Julia Drube, Carsten Hoffmann, Rob Leurs, Michel Bouvier, Meinrad P. Gawaz, An Hendrix, Etienne Moussay, Martine J. Smit, Jérôme Paggetti, Martyna Szpakowska, Andy Chevigné

## Abstract

Atypical chemokine receptors (ACKRs) are non-signaling GPCRs that regulate ligand availability, with ACKR3 functioning as a dual scavenger of chemokines and opioid peptides. Here, we demonstrate that following ligand stimulation, besides the canonical internalization, ACKR3 is released on extracellular vesicles (EVs). ACKR3 was also found on EVs released under basal conditions, although to a lesser extent. These observations were confirmed across multiple cellular contexts, including endogenous systems. Mechanistically, basal and ligand-induced EV release are independent of GRKs and β-arrestin but each relies on distinct trafficking routes and C-terminal determinants. Ligand-induced EV release is associated with plasma membrane localization and receptor recycling pathways. In contrast, basal EV release is governed by intracellular sorting processes and influenced by receptor ubiquitination and RAMP3. Functionally, EV-associated ACKR3 retains high-affinity ligand binding, enabling sequestration of CXCL12 and opioid peptides and thereby attenuating their signaling through CXCR4 and MOR. We also show that the release on EVs, in particular under basal conditions, is observed for other receptors such as KOR, CXCR4 and several ACKRs. Collectively, these findings establish EVs as regulators in chemokine and opioid systems and as a previously underappreciated dimension of ACKR3 and more broadly GPCR biology.

## 1 Introduction

Chemokine receptors are members of the G protein-coupled receptor (GPCR) family and mediate crucial homeostatic and inflammatory processes. They bind small chemotactic cytokines (chemokines) and induce G protein-mediated intracellular signaling ultimately leading to cell migration and proliferation. However, five atypical chemokine receptors (ACKR1–5) do not abide by this paradigm and have garnered increasing interest over the past years^1–3^. Indeed, these receptors are unable to trigger the activation of G proteins and subsequent signaling cascades upon ligand stimulation, yet they play important regulatory roles, including controlling ligand availability to shape chemokine gradients^1,4–7^.

ACKR3, formerly CXCR7, shares its two chemokine ligands CXCL12 and CXCL11 with the classical receptors CXCR4 and CXCR3, respectively. It plays a vital role in early cardiac and neuronal development ^8,9^, platelet activation^10^ as well as in the migration and homing of hematopoietic stem and progenitor cells^11–13^. The CXCL12–CXCR4–ACKR3 axis is heavily involved in the proliferation, migration and invasion of several cancer types, including breast cancer and leukemia^14–20^ and has shown promise as a target for anti-cancer and autoimmune therapies as well as several cardiovascular pathologies^18,21–25^. In addition, several non-chemokine endogenous peptides were recently described as ACKR3 agonists, including the proadrenomedullin N-terminal 20 peptides (PAMPs)^26^ and various opioid peptides especially from the enkephalin and dynorphin families^27^. Ligands triggering the phosphorylation of ACKR3 C tail, followed by β-arrestin recruitment and receptor–ligand endocytosis, are considered as agonists despite their inability to activate G proteins. While ACKR3 is substantially present intracellularly under basal conditions, it constitutively cycles from endosomal compartments to the plasma membrane, although the exact mechanism behind this shuttling has yet to be elucidated^28^. Several studies have linked ligand-free internalization to the C terminus of ACKR3^29,30^ and its basal ubiquitination^30^, while a more recent report found that it occurs in the absence of G protein-coupled receptor kinases (GRKs) and β-arrestins alike^31^. On the other hand, ligand-induced internalization of ACKR3 and subsequent ligand degradation were shown to rely on GRK phosphorylation and not require β-arrestins^31–35^. Additionally, ACKR3 ligand-induced trafficking and recycling has been suggested to be influenced by the presence of receptor activity-modifying proteins (RAMPs), especially RAMP3^36–38^.

Considerable research efforts have been devoted to elucidating the trafficking dynamics of ACKRs, and GPCRs in general. Recent evidence suggests a more intricate process than the traditionally reported cycling between the plasma membrane and internal compartments (*i.e*. recycling endosomes and degradative lysosomes)^31^. Moreover, several studies have also shown the shuttling of GPCRs in extracellular vesicles (EVs) and point to these as key mediators of intercellular communication^39–41^. EVs are bilayered lipidic structures that vary in size from 30 nm to 1000 nm and originate from endosomal compartments and/or the plasma membrane. They act as crucial cargo holders for a wide array of biomolecules including nucleic acids, proteins, and lipids and have essential functions in both physiological and pathological conditions^42,43^.

Importantly, many roles have been attributed to GPCR-harboring EVs, including specific inter-cellular receptor transfer and communication, the modulation of cell migration and even remote scavenging of GPCR ligands^40,44–49^.

In this study, using a panel of GPCR assays and EV characterization techniques compliant with the Minimal Information for Studies of Extracellular Vesicles (MISEV2023^50^) guidelines, we show that ACKR3 is released in EVs under basal conditions and in response to ligand stimulation, together with tetraspanin markers. These exports occur independently of β-arrestins and GRKs and are differentially linked to the receptor trafficking/recycling pathways, ubiquitination and interaction patterns with RAMPs. Profiling the 23 classical and atypical chemokine receptors reveals that ligand-induced EV release is unique to ACKR3 within this receptor family and is observed across multiple cellular contexts, including cells endogenously expressing the receptor such as platelets. In contrast, basal release also occurs for other receptors, notably ACKRs and can be detected in cancer cell lines, patient-derived samples and in vivo in a mouse model. We further demonstrate that ACKR3-containing vesicles, produced under both conditions bind and sequester the chemokine CXCL12 and the opioid peptide BAM22, thereby limiting their availability for the canonical signaling receptors CXCR4 and MOR, respectively. These findings extend ACKR3 function beyond the cell surface and identify EVs as regulators of chemokine and opioid peptide signaling and highlight a possible broader and underappreciated role of EVs in GPCR biology.

## 2 Results

### 2.1 Chemokines and opioid peptides trigger both ACKR3 internalization and release in extracellular vesicles

ACKR3 is recognized as a receptor that constitutively cycles from endosomal compartments to the plasma membrane, without requiring β-arrestins^31,32,34,51^. Many studies unitedly report that ACKR3 is rapidly internalized following the activation by chemokines or short peptides^28–30,34^.

Using flow cytometry and a bystander NanoBRET assay, we confirmed that the binding of CXCL12 or the proenkephalin-derived peptide BAM22 reduces ACKR3 presence at the cell surface compared to basal conditions or stimulation with the irrelevant chemokine CCL5 (**Fig. 1a**). However, the use of a novel, sensitive alternative method to assess the presence of the receptor at the plasma membrane allowed us to uncover a process parallel to receptor internalization. Indeed, the NanoGlo HiBiT extracellular assay, which quantifies the presence of HiBiT-tagged receptor through Nanoluciferase complementation with membrane-impermeable LgBiT protein fragment, revealed a seeming four-fold increase of receptor presence upon CXCL12 stimulation (**Fig. 1b–c, left**). Interestingly, we found a similar increase of ACKR3 in the isolated cell supernatant following CXCL12 or BAM22 stimulation (**Fig. 1c, right**). We hypothesized that this observation resulted from a release of ACKR3 in EVs, which are typically overlooked in approaches quantifying the receptor only at the plasma membrane with specific markers (e.g. bystander BRET) or requiring washing steps during which these EVs are discarded (e.g. flow cytometry, ELISA). This was further supported by quantitative analysis of particle concentration in the supernatant of CXCL12-stimulated cells, demonstrating a similar increase to that observed in the HiBiT assay (**Fig. 1d**).

**Figure 1:**
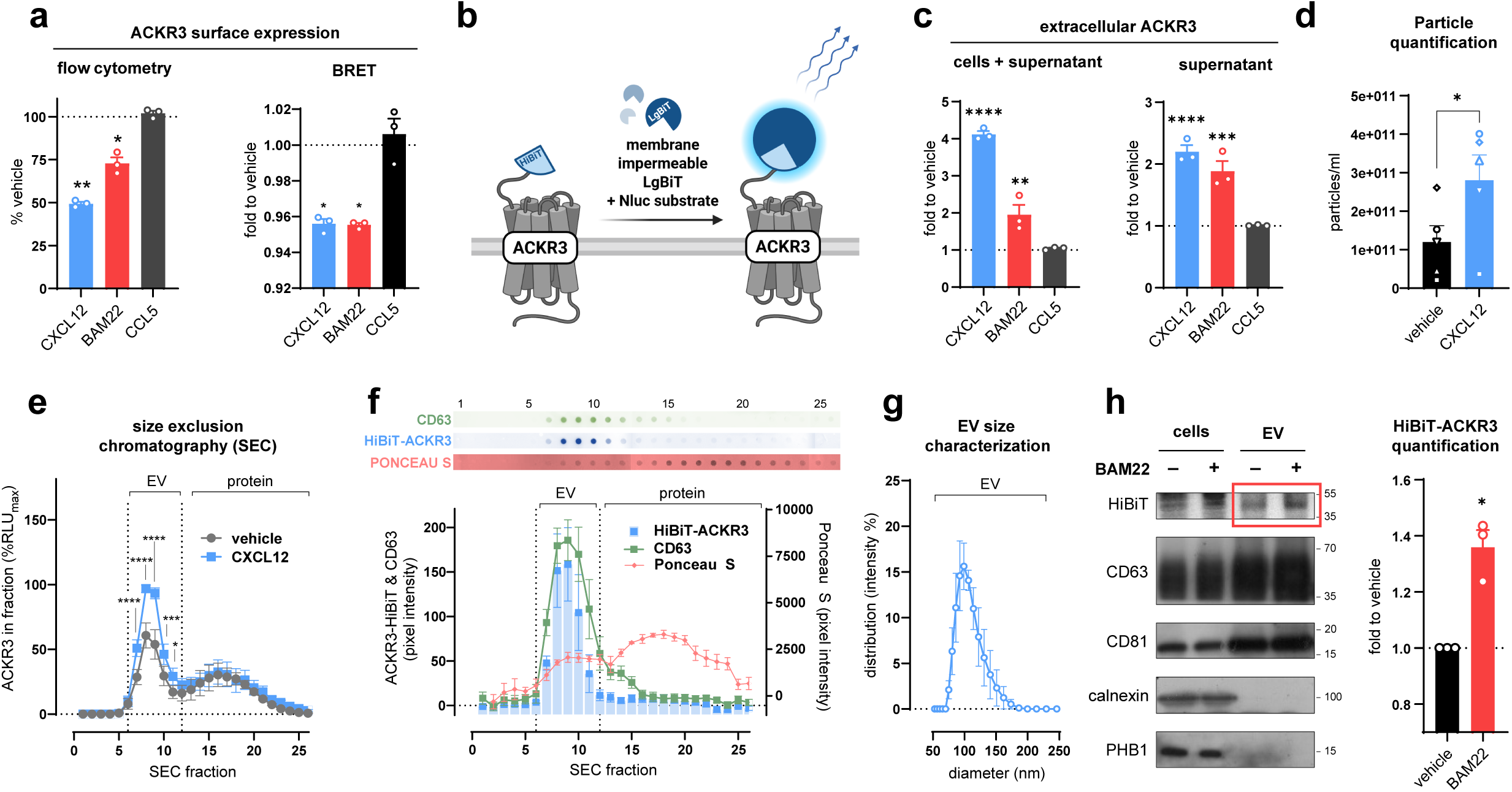
ACKR3 is released on extracellular vesicles upon ligand stimulation. **(a)** ACKR3 internalization after stimulation with CXCL12, CCL5 (100 nM) or BAM22 (1 µM) monitored by flow cytometry on HEK293T cells stably expressing HA-ACKR3 (*left*) or bystander NanoBRET (*right*) on transiently transfected HEK293T cells using a Lyn-derived plasma membrane anchor. **(b)** Schematic representation of the HiBiT complementation assay to monitor receptor presence at the cell surface used in c: addition of membrane impermeable LgBiT and substrate to HiBiT-ACKR3-expressing cells following ligand stimulation. **(c)** Detection of HiBiT-tagged receptors through Nluc complementation prior (*left*) and after (*right*) supernatant isolation through centrifugation. **(d)** Cumulative count of particles between 50 and 200 nm measured by dynamic light scattering (DLS) in supernatants isolated through centrifugation from HEK293T cells transiently expressing HiBiT-ACKR3 and stimulated or not with CXCL12 (100 nM). **(e)** Size exclusion chromatography (SEC) analysis of the isolated supernatants used in (d) followed by HiBiT complementation in the different fractions. **(f)** Representative dot blot images of each SEC fraction (*top*) and quantification across fractions (*bottom*) of HiBiT-ACKR3, CD63 and total protein (Ponceau S) from supernatants stimulated with CXCL12 (100 nM). **(g**) Size distribution of pooled SEC-derived EV fractions from supernatants of CXCL12-stimulated cells measured by DLS. Each population is weighted by its relative contribution to the total scattered light signal. **(h)** Representative western blot of the lysates of HEK293T cells stably expressing HiBiT-ACKR3 stimulated with BAM22 (250 nM) or vehicle and EVs isolated by ultracentrifugation from the supernatant (*left*). Blots were probed for HiBiT, the EV markers CD63 and CD81, and the purity controls calnexin and PHB1. Quantification of ACKR3-HiBiT signal from three independent EV purifications as shown on the left *(right)*. Results are presented as mean ± S.E.M of at least three independent experiments (n ≥ 3). *p < 0.05, **p < 0.01, ***p < 0.001, ****p < 0.0001 by lognormal and normal ordinary one-way ANOVA with Dunnet’s correction (a, c respectively), by two-tailed ratio paired t-test (d, h), by two-way repeated measures ANOVA with Šídák’s multiple comparisons test (e). While normalized data are displayed, statistical analyses were performed on raw values.

To isolate EV population, we applied size exclusion chromatography (SEC) as our first purification method. SEC effectively separates EVs from unfolded, degraded, or aggregated free proteins present in the supernatant^52^. It revealed a significant increase of ACKR3 following CXCL12 stimulation within fractions corresponding to EVs (fractions 6–12) and not in the soluble protein fractions (**Fig. 1e**). This was supported by the enrichment of the tetraspanin EV marker CD63 in these fractions (**Fig. 1f**) and a particle size distribution of 75**-**150 nm consistent with that of EVs (**Fig. 1g**). As an orthogonal validation approach, we isolated EV-enriched fractions from culture supernatants of untreated and BAM22-treated cells using differential ultracentrifugation followed by western blot analysis, which confirmed an increase in intact HiBiT-tagged ACKR3 on EVs following stimulation. Consistently, these fractions were enriched in the tetraspanin markers CD81 and CD63, while lacking intracellular organelle markers calnexin and PHB1 (**Fig. 1h**).

### 2.2 A selective ligand-induced extracellular vesicle release yields a pool of functional ACKR3

Next, we further characterized this ligand-induced release of ACKR3-containing EVs. We showed that this phenomenon was ligand concentration-dependent with potencies in the nanomolar range (**Fig. 2a**) and followed a trend of rapid initial increase reaching a plateau by one hour (**Fig. 2b**). Furthermore, increased ACKR3 measured in the supernatant correlated with the global expression levels of ACKR3 (**Fig. 2c**). We also found that ACKR3 release was not restricted to a single cellular context as it was observed in the supernatant of several other cell types, including U87-MG (malignant glioma), HeLa (cervical cancer) and MCF-7 (breast cancer) transiently expressing ACKR3 (**Fig. 2d)**. Similar observations of extracellular ACKR3 increase were made for the murine orthologue (**Fig. 2e**).

**Figure 2:**
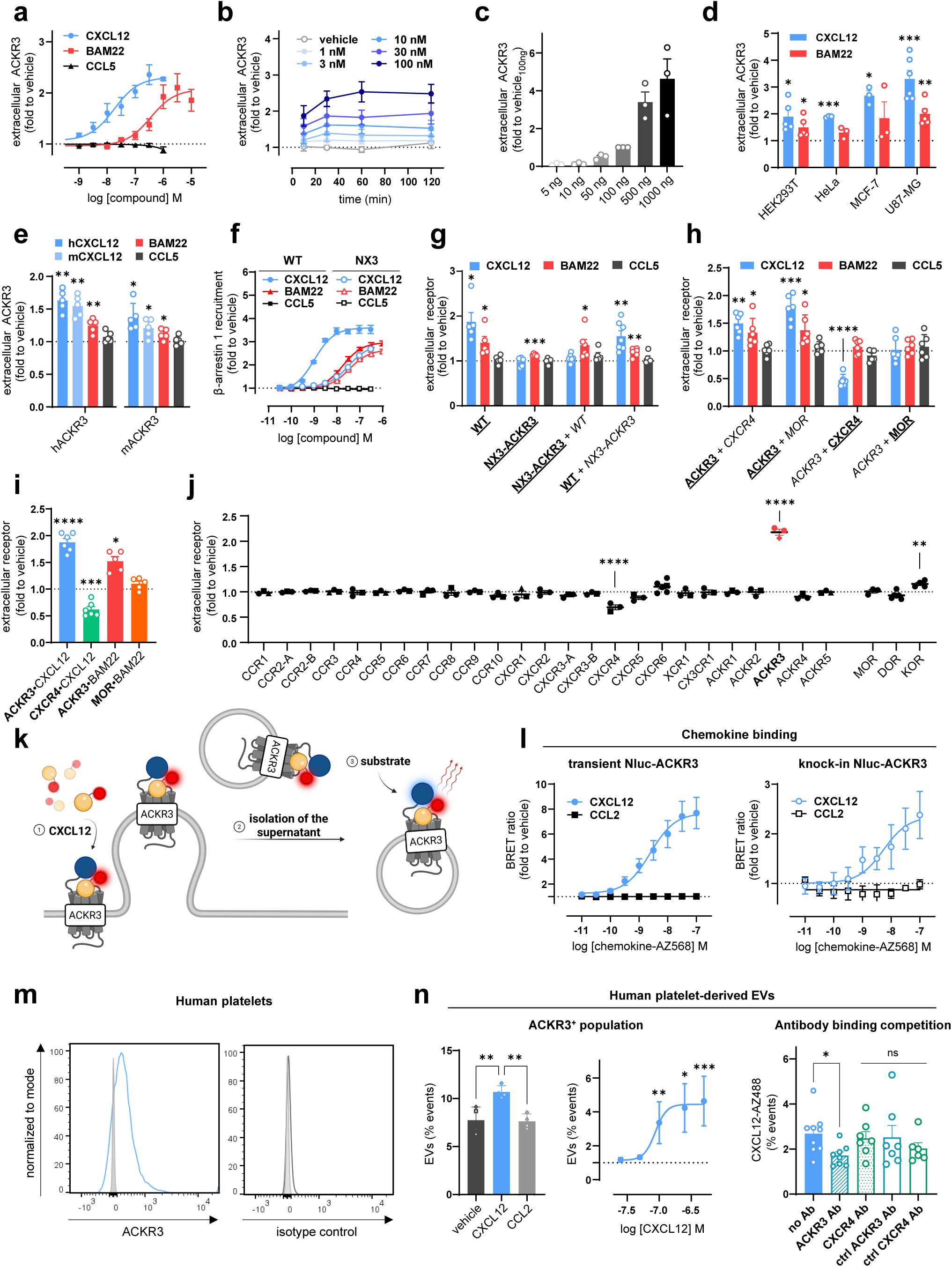
Ligand-induced release in EVs is specific to ACKR3. **(a)** Concentration–response relationship of release of ACKR3-harboring vesicles treated with CXCL12, BAM22 or CCL5 as a negative control monitored by Nluc complementation of HiBiT-tagged ACKR3. **(b)** Kinetics of CXCL12-induced release of ACKR3-harboring EVs following stimulation with different concentrations of CXCL12. **(c)** Dependency of ACKR3 release on the overall ACKR3 expression level. Increasing amounts of plasmid encoding HiBiT-ACKR3 were transfected (per 1 μg total DNA transfected). Cells were treated with CXCL12 and signal was normalized to total receptor expression and to the 100 ng stimulated condition. **(d)** ACKR3 presence in the supernatant of HEK293T, HeLa, MCF-7 and U87 MG cells expressing HiBiT-ACKR3 and treated with CXCL12 or BAM22. **(e)** Presence of human and mouse ACKR3 quantified in the supernatant of transfected HEK293T cells treated with human or mouse CXCL12, BAM22 and CCL5. **(f)** β-arrestin 1 recruitment to ACKR3 and CXCR3-N terminus ACKR3 (NX3-ACKR3) chimera induced by CXCL12, BAM22 or CCL5 determined by NanoBiT. **(g)** Presence of HiBiT-tagged (*bold, underlined*) WT or NX3-ACKR3 chimera in the supernatant when co-expressed (1:1 ratio) with and without untagged WT or NX3-ACKR3 chimera (*italic*) and stimulated with CXCL12, BAM22 or CCL5. **(h)** Presence of HiBiT-tagged (*bold*) ACKR3, CXCR4 or MOR in the supernatant when expressed alone or co-expressed (1:1 ratio) with untagged ACKR3, CXCR4 or MOR (*italic*) and stimulated with CXCL12, CCL5 or BAM22. **(i)** HiBiT-tagged receptors CXCR4, ACKR3 and MOR presence in the supernatant after stimulation with CXCL12 or BAM22. **(j)** Ligand-induced extracellular release of HiBiT-tagged receptors. Ligand–receptor pairs are listed in Supplementary Table 1. **(k)** Schematic representation of the NanoBRET assay used to measure sustained chemokine binding to ACKR3-harboring vesicles in the isolated supernatant: (1) stimulation of Nluc-tagged ACKR3 with fluorescently labelled chemokine; (2) isolation of the supernatant (3) addition of substrate and NanoBRET measurement. **(l)** Binding of fluorescently labelled CXCL12 or CCL2 to ACKR3-harboring EVs measured by NanoBRET in the supernatant harvested after stimulation of HeLa cells transiently expressing ACKR3 N-terminally tagged with Nluc (*left*) or CRISPR-edited knock-in HeLa cells expressing endogenous level of Nluc-ACKR3 fusion protein (*right*). **(m)** Representative flow cytometry histograms showing ACKR3 staining (left) and isotype control (right) on isolated platelets from a healthy donor. Unstained cells are shown in grey. **(n)** *(left)* Quantification of EV release under shear stress from human platelets stimulated with CXCL12 (250 nM), CCL2 (250 nM) or left unstimulated. *(middle)* Concentration–response curve of CXCL12-induced EV release from human platelets. *(right)* CXCL12-AZ488 ligand binding on human platelets alone or co-incubated with an anti-ACKR3 monoclonal antibody. All experiments were performed in HEK293T cells unless otherwise stated. CXCL12, CCL2 and CCL5 were used at 100 nM and opioid peptides at 1 µM. Results are presented as mean ± S.E.M of at least three independent experiments (n ≥ 3). *p < 0.05, **p < 0.01, ***p < 0.001, ****p < 0.0001 by one sample t-test comparing a normalized value of 1 (d–e, g–h), two-way repeated measures ANOVA with Šídák’s multiple comparisons test (i–j) or by one-way ANOVA with Tukey’s or Dunnet’s correction (n left, n middle/right respectively).

To better understand the mechanism driving the ligand-induced release, we designed a chimeric ACKR3 harboring the N terminus of CXCR3 (NX3-ACKR3). The agonist activity in β-arrestin recruitment of CXCL12, but not BAM22, was markedly reduced towards this receptor **(Fig. 2f)**. Accordingly, CXCL12 failed to induce release of NX3-ACKR3, whereas BAM22 retained this ability, suggesting that indeed specific ligand-bound conformations of the receptor are required for this process (**Fig. 2g**). Importantly, the co-expression of untagged WT ACKR3 did not lead to a co-release of NX3-ACKR3 following CXCL12 stimulation, further supporting that direct ligand binding is necessary. We also investigated whether other receptors could be co-exported with ACKR3. Using co-expression of HiBiT-tagged CXCR4 and MOR, we found that the stimulation of ACKR3 did not lead to their co-release, pointing towards the selective sorting of ACKR3 onto EVs rather than unspecific membrane budding (**Fig. 2h**).

We further assessed whether EV release may be inherent to the ligands used and hence investigated the potential release of receptors with overlapping ligand profiles. No increase was observed for the classical chemokine receptor CXCR4, or the opioid receptor MOR in response to CXCL12 and BAM22, respectively (**Fig. 2i**). We also wondered if the release of receptors in the supernatant upon ligand binding was unique to ACKR3 or if any other member within the chemokine family behaved similarly. We therefore screened all human chemokine receptors using the extracellular HiBiT assay, which revealed that ACKR3 was the only chemokine receptor to be significantly increased in the supernatant in response to a ligand (**Fig. 2j**). While changes were observed for other receptors, including CXCR4, further SEC analysis demonstrated that these originated from the soluble protein fraction rather than EV-associated fractions (**Supplementary Fig. 2**). Additionally, the opioid receptor KOR exhibited a slight increase following stimulation, which is consistent with previous reports suggesting its packaging in EVs^53,54^.

We next evaluated whether the ligand still interacts with EV-associated ACKR3 once released. To assess the presence and stability of ligand–receptor complexes in the supernatant, we applied a NanoBRET binding assay using cells expressing ACKR3 N-terminally fused to the full Nanoluciferase (Nluc) (**Fig. 2k)**. We first measured NanoBRET in the supernatant of ACKR3-expressing cells stimulated with fluorescently labelled CXCL12. Using HeLa cells overexpressing Nluc-ACKR3, we found a concentration-dependent increase in BRET signal in the supernatant indicating that the receptor is properly folded on the EV surface and retains its ability to bind CXCL12 with nanomolar potency, consistent with the reported affinity of the chemokine for the receptor^28^ (**Fig. 2l, left**). We observed a similar ligand-induced release of ACKR3-EVs and a strong and stable interaction with CXCL12 using HeLa knock-in cells CRISPR-edited to express the Nluc-tagged ACKR3 (**Fig. 2l, right**). These results suggest that in addition to intracellular scavenging via internalization of receptor–chemokine complexes, sequestration of chemokines may occur in parallel through binding to released ACKR3-bearing EVs.

Supporting the physiological relevance of these findings, we showed that ligand-induced EV release and chemokine capture were also observed in human platelets, where ACKR3 is endogenously expressed (**Fig. 2m)**^10,55,56^. Indeed, under shear stress conditions, stimulation of platelets with CXCL12 triggered the release of ACKR3-containing EVs in a concentration-dependent manner (**Fig. 2n, left and middle**). Furthermore, we showed that these EVs can efficiently sequester the chemokine at their surface which can, in turn, be displaced by an anti-ACKR3 monoclonal antibody, excluding CXCR4 engagement and overall supporting observations made in cell line models (**Fig. 2n, right**).

Taken together, these data highlight a rapid and significant release of ACKR3 in the extracellular space via EVs, which is ligand- and concentration-dependent and unique within the chemokine receptor family. While ACKR3 is internalized from the plasma membrane in response to ligand, the finding that it is also present on EVs sheds light on a previously overlooked compartment, potentially expanding the scope of this receptor’s biological activity.

### 2.3 ACKR3 is also released on extracellular vesicles under basal conditions

Considering the substantial proportion of ACKR3-containing EVs observed in unstimulated cells (**Fig. 1e**), we also explored the extent of this release and its possible functional relevance.

Analysis revealed a continuous release of ACKR3-positive EVs over time under basal conditions (**Fig. 3a**). SEC fractionation of conditioned medium confirmed the presence of ACKR3 in EV-enriched fractions as evidenced by the co-enrichment of CD63 (**Fig. 3b–c**). As an independent validation approach, western blot analysis of EV-enriched preparation purified by ultracentrifugation further confirmed the presence of intact HiBiT-tagged ACKR3 alongside canonical EV markers CD63 and CD81, while particle size analysis indicated a vesicle population ranging from 75 to 150 nm (**Fig. 3d–e**). Purified EV populations were analysed using the recently developed single-EV high-sensitivity nanoscale fluorescence-activated technology (**Fig. 3f**) as well as bulk-EV flow cytometry (**Fig. 3g, Supplementary Fig. 3**) and single-EV spectral flow cytometry (**Fig. 3h–i**). Using a conformational anti-ACKR3 antibody instead of HiBiT detection, these three independent approaches confirmed the presence of intact receptor on EVs together with tetraspanin markers CD9, CD63 and CD81. Interestingly, single-EV analysis revealed that 10% of the total EV population was positive for ACKR3 and that all ACKR3-positive EVs contained both CD63 and CD81 (**Fig. 3h–i**). Finally, super-resolution fluorescence imaging (direct stochastic optical reconstruction microscopy, dSTORM) of single EV particles confirmed the presence of ACKR3 at the surface of the vesicles together with the tetraspanin markers (**Fig. 3j**).

**Figure 3:**
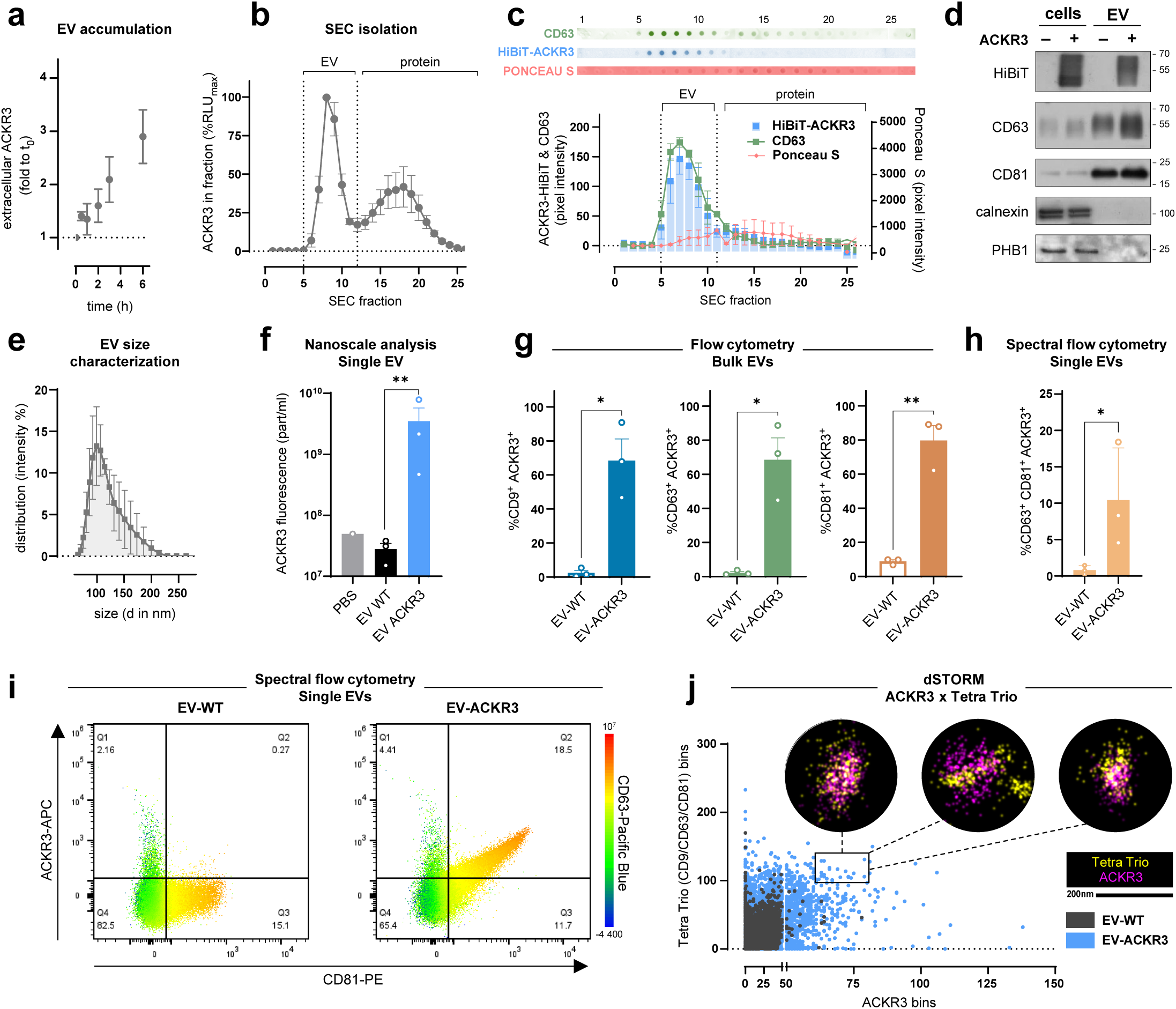
ACKR3-harboring vesicles are released under basal conditions. **(a)** Basal release of ACKR3-harboring vesicles by HEK293T cells stably expressing HiBiT-ACKR3 and measured by HiBiT complementation in supernatant collected over six hours. **(b)** SEC fractionation of supernatants of HEK293T cells transiently expressing HiBiT-ACKR3 collected after 30 minutes, revealed by complementation with LgBiT. **(c)** Representative dot blot images of each SEC fraction (*top*) and quantification of HiBiT-ACKR3, CD63 and total protein (Ponceau S) signals across fractions (*bottom*). **(d)** Representative western blot of the lysates of parental HEK293T cells (ACKR3-negative (-)) and stably expressing HiBiT-ACKR3 (ACKR3-positive (+)) and EVs isolated by ultracentrifugation from the supernatant. Blots were probed for HiBiT, the EV markers CD63 and CD81, and the purity controls calnexin and PHB1. **(e)** Size distribution of EVs isolated by ultracentrifugation from cells stably expressing HiBiT-ACKR3, measured by DLS. **(f)** Quantification of ACKR3-positive EVs by fluorescence-based nanoparticle tracking (Hekat) in PBS control, EVs originating from parental HEK293T (EV-WT), or from ACKR3-expressing HEK293T cells (EV-ACKR3). **(g)** Flow cytometry analysis of bead-captured EV-WT and EV-ACKR3, showing the proportion of CD9L, CD63L, or CD81L events that are ACKR3L. **(h)** Proportion of CD63LCD81LACKR3L in single-EV resolution in EV-WT and EV-ACKR3 preparations, assessed by spectral flow cytometry. **(i)** Representative spectral flow cytometry dot plots of single EVs from EV-WT and EV-ACKR3 preparations, showing ACKR3 signal (y-axis, detected with 8F11-M16 antibody) against CD81 signal (x-axis), with CD63 intensity represented by colour scale. **(j)** dSTORM super-resolution analysis of EVs isolated by ultracentrifugation from wild-type (EV-WT) or HiBiT-ACKR3-expressing (EV-ACKR3) HEK293T cells, labelled with anti-ACKR3 and anti-tetraspanin (Tetra Trio: CD9/CD63/CD81) antibodies. Scatter plot shows the number of Tetra Trio bins versus ACKR3 bins per individual EV. *Inset:* representative dSTORM images of EVs from the boxed region. Results are presented as mean ± S.E.M of at least three independent experiments (n ≥ 3). *p < 0.05, **p < 0.01 by lognormal one-way ANOVA (f), or one-tailed paired or unpaired t-test (g, h respectively).

### 2.4 ACKR3-harboring vesicles released under basal conditions are capable of ligand neutralization

We subsequently wondered whether EVs released under basal conditions could also bind and sequester ACKR3 ligands in the extracellular space. Indeed, we could detect a concentration-dependent NanoBRET signal upon addition of fluorescently labelled CXCL12 to the supernatant isolated from HeLa cells transiently or endogenously expressing Nluc-ACKR3, revealing chemokine binding in the subnanomolar potency range (**Fig. 4a and Supplementary Fig. 4**). Post-SEC NanoBRET analysis confirmed that this binding was exclusively detected in the EV fractions, confirming the integrity and functionality of ACKR3 displayed at the surface of the vesicles (**Fig. 4b–c**). Screening all human chemokine receptors revealed that, in addition to ACKR3, all other four ACKRs as well as CXCR4 are exported extracellularly under basal conditions (**Fig. 4d**). Importantly, further post-SEC NanoBRET analysis showed that also these receptors are functional, retaining their ability to bind their respective chemokines in the EV fractions exclusively (**Fig. 4d inset**).

**Figure 4:**
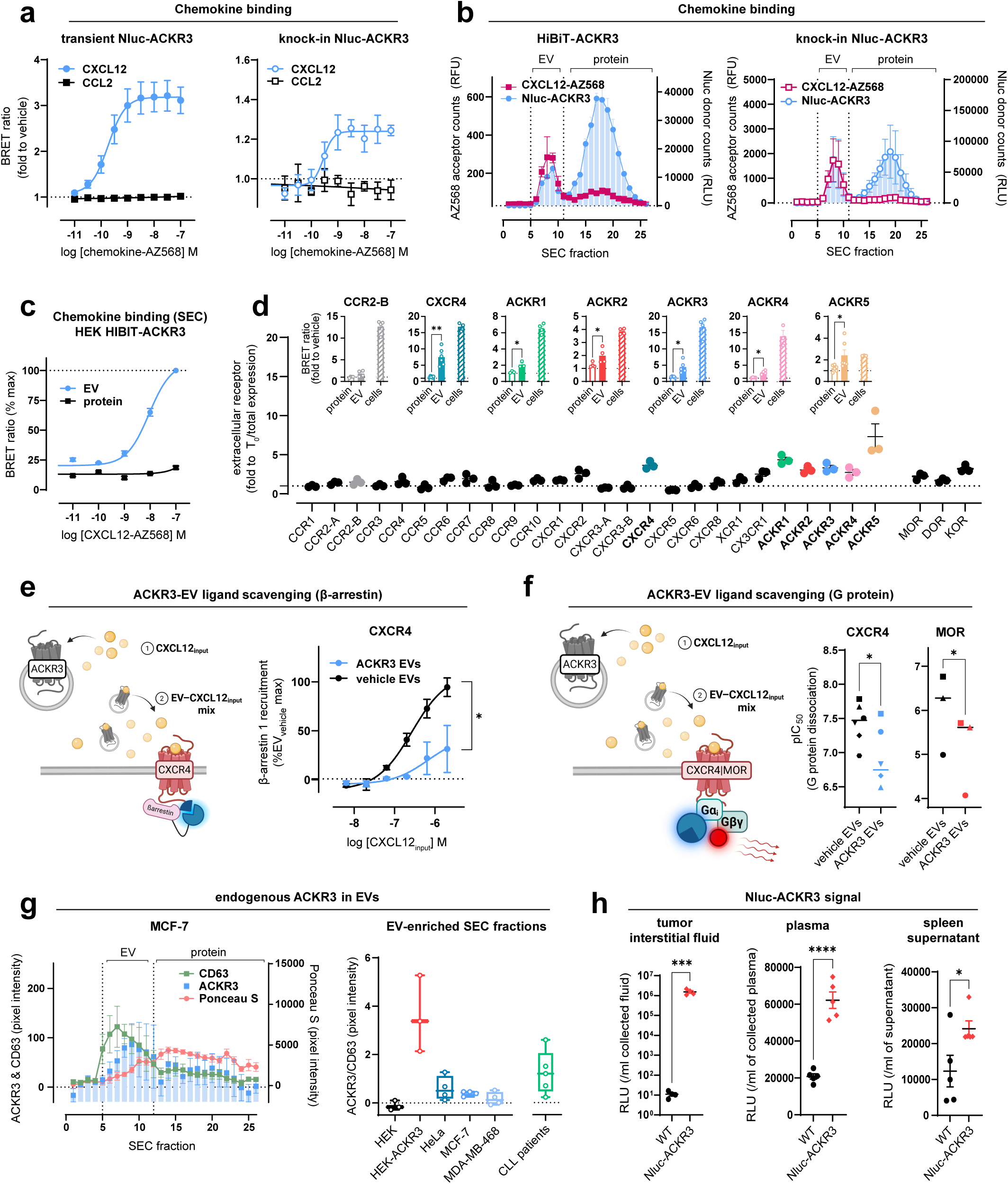
ACKR3-harboring vesicles released under basal conditions are capable of ligand neutralization. **(a)** Concentration-dependent binding of CXCL12- and CCL2-AZ568 to ACKR3-harboring vesicles released under basal conditions from HeLa cells transiently expressing Nluc-ACKR3 (*left*) or CRISPR-edited to express Nluc-ACKR3 (*right*). **(b)** NanoBRET analysis of individual SEC fractions of supernatants from HEK293T cells expressing HiBiT-ACKR3 (left) or CRISPR-edited Nluc-ACKR3 HeLa cells (*right*) incubated with CXCL12-AZ568 (200 nM). NanoBRET donor (Nluc luminescence, via LgBiT complementation) and acceptor (AZ568 fluorescence) signals are shown separately across SEC fractions. **(c)** NanoBRET binding curves for CXCL12-AZ568 applied to pooled SEC fractions corresponding to EV-enriched or soluble protein-enriched fractions from HiBiT-ACKR3-expressing HEK293T cells. **(d)** Constitutive, unstimulated release of HiBiT-tagged chemokine receptors over six hours, normalized to release at T_0_. *Insets*: NanoBRET ligand binding signal across SEC-separated pooled EV- and protein-enriched fractions, as well as intact cells for CCR2-B (CCL2), CXCR4 (CXCL12), ACKR1 (CCL2), ACKR2 (CCL2), ACKR3 (CXCL12), ACKR4 (CCL19), ACKR5 (CXCL12). All chemokines were labeled with AZ568 and used at 100 nM. **(e)** Schematic representation of the β-arrestin ligand scavenging assay (*left*): (1) co-incubation of concentrated conditioned medium of ACKR3-expressing cells (ACKR3 EVs) or untransfected cells (vehicle EVs) with CXCL12 at concentrations referred to as CXCL12_input_, (2) CXCL12–EV mix was diluted 20-fold and used to induce β-arrestin recruitment to CXCR4 monitored by NanoBiT on transfected HEK293T. (*right*) β-arrestin 1 recruitment to CXCR4. x-axis corresponds to the CXCL12 concentrations added to ACKR3 and vehicle EVs before a 20-fold dilution on the receiving signaling cells. **(f)** Schematic representation of the G protein dissociation ligand scavenging assay (*left*): (1) co-incubation of concentrated conditioned medium of ACKR3-expressing cells (ACKR3 EVs) or untransfected cells (vehicle EVs) with CXCL12 or BAM22 at concentrations from which the pIC_50_ values were calculated, (2) ligand–EV mix used to induce Gα_i_ dissociation in HEK293T cells expressing CXCR4 or MOR monitored by NanoBRET. (*right*): pIC_50_ values were calculated based on the ligand concentrations added to ACKR3 and vehicle EVs before a 20-fold dilution on the receiving signaling cells. Each symbol represents a biological replicate. **(g)** Quantification of ACKR3, CD63, and total protein dot blot signal across SEC fractions of conditioned MCF-7 media (*left*). ACKR3/CD63 signal ratio in pooled EV-enriched fractions (6–12) from WT HEK293T (HEK) or HEK293T transiently expressing HiBiT-ACKR3 (HEK-ACKR3), three cell lines (HeLa, MCF-7, MDA-MB-468) endogenously expressing ACKR3 and CLL patient samples (n = 5 donors). Each point is an independent biological replicate. **(h)** Nluc signal measured in tumor interstitial fluid, blood plasma, and spleen supernatant from mice injected with wild-type (WT) or CRISPR knock-in Nluc-ACKR3 (Nluc-ACKR3) HeLa cells (n = 5 mice per group). If not stated otherwise, results are presented as mean ± S.E.M of at least three independent experiments (n ≥ 3). *p < 0.05, ***p < 0.001, by two-tailed paired t-test (d-inset, f), two-sample paired Wilcoxon test applied to paired points at different concentrations (e) and two-tailed unpaired Mann-Whitney test (h).

We next asked whether ACKR3-harboring EVs released under basal conditions are capable of sequestering chemokines and opioid peptides to reduce signaling through classical receptors such as CXCR4 and MOR. Incubation of CXCL12 and BAM22 with supernatant isolated from ACKR3-expressing cells significantly reduced their ability to activate their respective cognate signaling receptor, as shown in β-arrestin recruitment **(Fig. 4e)** and in G protein activation assays **(Fig. 4f)**. To further explore the potential physiological relevance, we also confirmed that the basal release of ACKR3-harboring EVs can be detected using anti-ACKR3 antibody in the EV fractions of supernatant purified from various cancer cell lines endogenously expressing ACKR3, including HeLa, MCF7 and MBA-MB-468 as well as from cells of chronic lymphocytic leukemia (CLL) patients **(Fig. 4g).** Finally, in mice bearing localized subcutaneous tumors derived from Nluc-ACKR3 knock-in HeLa cells, Nluc signal could be detected not only in the isolated tumor interstitial fluid but also in the plasma and purified spleen supernatant, suggesting the presence of ACKR3-containing vesicles at distant sites and potential remote regulation of ligand availability **(Fig. 4h)**.

Altogether, these findings indicate that ACKR3-positive EVs released under basal conditions can bind and sequester ligands, revealing a potential mechanism for the remote regulation of ligand bioavailability. Notably, the extracellular export of functional receptors in the absence of stimulation does not appear to be limited to ACKR3, as several other chemokine receptors are also detected on EVs under basal conditions, suggesting that release of receptor-containing EVs may represent a broader and previously underappreciated feature of GPCR biology.

### 2.5 ACKR3-harboring extracellular vesicles released under ligand-induced and basal conditions arise from different cellular compartments

We then investigated whether ACKR3 EVs released under ligand-induced and basal conditions share a common cellular origin or arise from different compartments, and how their release may relate to the atypical intracellular localization or trafficking dynamics of the receptor.

We found that particles released under each condition exhibited slightly different size distributions, consistent with the possibility that they originate from sorting pathways and/or cellular compartments (**Fig. 5a**).

**Figure 5:**
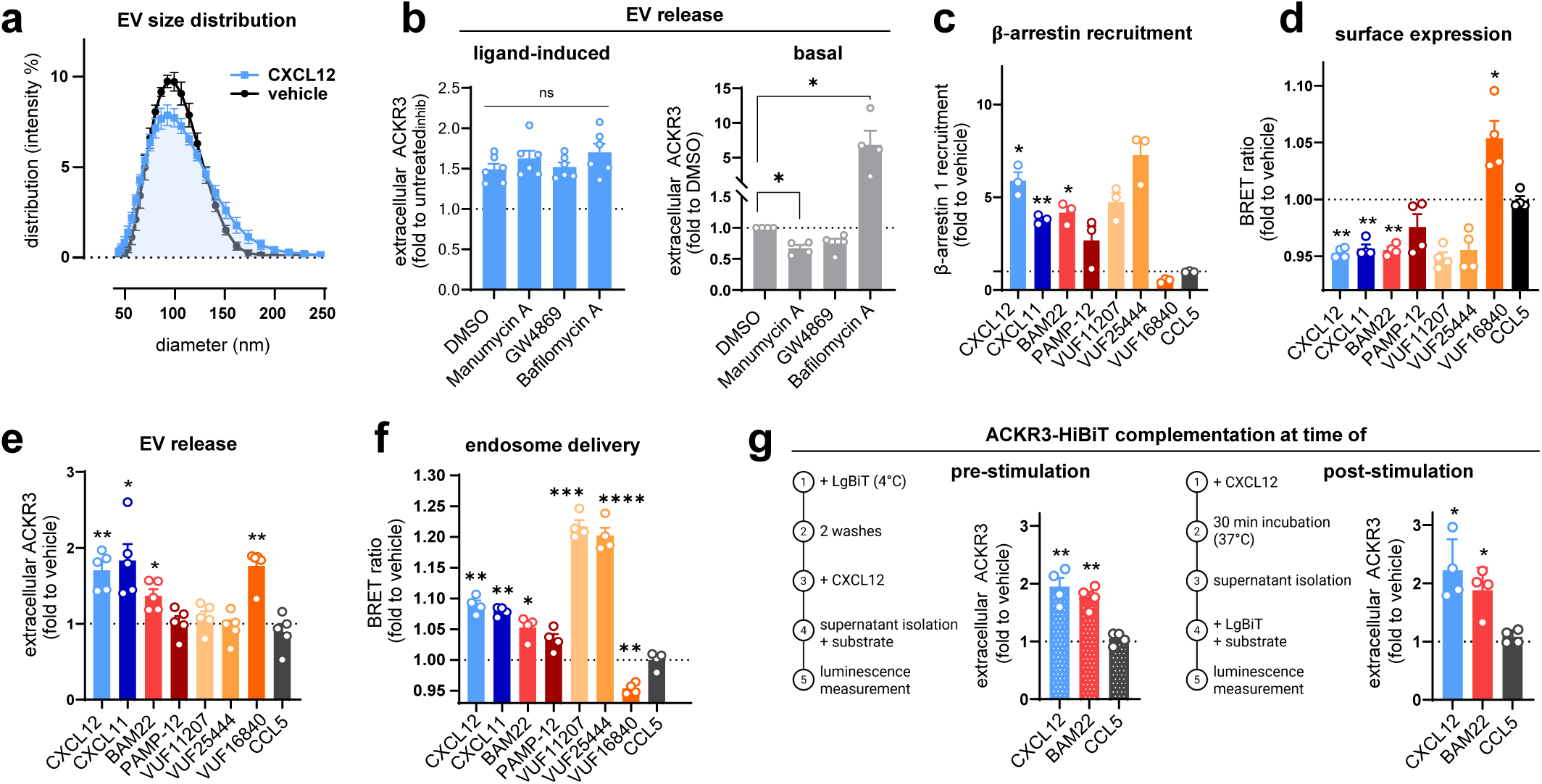
ACKR3-harboring vesicles released under ligand-induced and basal conditions arise from different cellular compartments. **(a)** Size distribution measured by DLS of supernatant isolated from HiBiT-ACKR3-expressing cells stimulated with CXCL12 or left unstimulated. **(b)** Quantification of ACKR3-harbouring vesicles released in the supernatant of HEK293T cells expressing HiBiT-ACKR3 in response to CXCL12 (100 nM) (*left*) or recovered under basal conditions (*right*), following overnight treatment with manumycin A (250 nM), GW4869 (250 nM) or bafilomycin A1 (250 nM). **(c–f)** Ligand-induced effect on **(c)** β-arrestin 1 recruitment to ACKR3 monitored by NanoBiT, **(d)** surface expression of ACKR3 monitored by bystander NanoBRET, **(e)** extracellular release of ACKR3-harboring vesicles monitored by HiBiT–LgBiT complementation in the isolated supernatant and **(f)** delivery to early endosomes monitored by bystander NanoBRET. **(g)** ACKR3-harboring vesicles released from the cell surface in response to CXCL12, CCL5 or BAM22 monitored using HiBiT–LgBiT complementation. Cells were pre-incubated with LgBiT prior stimulation to complement HiBiT-ACKR3 at the cell surface exclusively (*left*), or the usual set-up of stimulation, EV isolation and LgBiT complementation was applied (*right*) (see Supplementary Fig. 5 for further details). Chemokines CXCL12, CXCL11 and CCL5 were used at 100 nM and peptides BAM22, PAMP-12 and small molecules VUF11207, VUF25444 and VUF16840 were used at 1 µM. Results are presented as mean ± S.E.M of at least three independent experiments (n ≥ 3). *p < 0.05, **p < 0.01, by lognormal one-way ANOVA with Dunnett’s correction (b–g). While normalized data are displayed, statistical analyses were performed on raw values.

To explore this further, we used pharmacological modulators of EV biosynthesis and sorting pathways, which were described to decrease (manumycin A and GW4869) or increase (bafilomycin A1) the release of EVs of intracellular origin but not membrane-derived EVs^57–60^. Interestingly, we observed that while ligand-induced release was not affected by any of these inhibitors, the basal release of ACKR3-positive EVs was reduced by manumycin A and, to some extent, GW4869 and largely enhanced by bafilomycin A1, pointing to an intracellular origin **(Fig. 5b)**.

To gain further insight into the origin and mechanism of ligand-induced ACKR3-harboring EV-release, we next took advantage of the broad pharmacological diversity of ACKR3 ligands, including molecules known to perturb the basal cycling and localization of the receptor. Beyond CXCL12 and BAM22, we tested CXCL11, as well as the endogenous peptide agonist PAMP-12^26^ and several small molecules acting as full (VUF11207^61^), super (VUF25444^62^) or inverse agonists (VUF16840^63^). We first validated the respective pharmacological profiles of the ligands using a β-arrestin recruitment assay, considered as standard to monitor the activity of ACKR3 (**Fig. 5c**). We next demonstrated with a bystander NanoBRET assay that all endogenous and synthetic agonists were able to trigger receptor disappearance from the plasma membrane, except for the inverse agonist VUF16840, which induced an increase of surface ACKR3 levels (**Fig. 5d**)^64^.

We then analyzed the ability of these different modulators to trigger the release of ACKR3-harboring vesicles. We found that, much like CXCL12 and BAM22, CXCL11 induced robust ACKR3 release. Interestingly, small-molecule agonists such as VUF11207 or the superagonist VUF25444 were poor inducers of EV release (**Fig. 5e**), indicating that the ability to induce EV release does not directly correlate with ligand capacity to trigger β-arrestin recruitment. Moreover, VUF11207 and VUF25444 showed a superior ability to induce receptor endocytosis (**Fig. 5f**), as indicated by the stronger BRET signal with tagged FYVE in the endosomal compartment, pointing to a role for ligand-specific receptor engagment in directing the trafficking of ACKR3. Hence, the reduction in surface receptor measured with these ligands (**Fig. 5d**) is driven almost exclusively towards internalization and delivery to endosomes, and not to EV release. In contrast, treatment with the inverse agonist VUF16840 reduced basal ACKR3 trafficking through early endosomes, increased receptor accumulation at the plasma membrane, and enhanced the release of ACKR3-harboring EVs (**Fig. 5d–f**). Together, these findings suggest that ligand-induced changes in ACKR3 conformation and its interactions with scaffolding or retention proteins can differentially gate receptor entry into endocytic versus EV-sorting pathways.

To further establish if the receptors released upon stimulation originate directly from the cell surface or are mobilized from an intracellular pool, we incubated cells on ice during ACKR3-HiBiT complementation to exclusively track receptors present at the cell surface before stimulation (**Supplementary Fig. 5**). We found a similar increase of ACKR3 in the isolated supernatant as with the initial setup (**Fig. 1e**), indicating that a large part of the receptors released originate from a pool already present at the plasma membrane at the time of ligand stimulation (**Fig. 5g**).

### 2.6 Ligand-induced and basal release is independent of GRKs and β-arrestins, and differentially affected by receptor C terminus motifs and RAMP interactions

The activation and trafficking patterns of ACKR3 have been extensively studied and it is well established that ligand binding triggers phosphorylation of the intracellular regions of the receptor by G protein-coupled receptor kinases (GRKs). This leads to the recruitment of β-arrestins and eventually to the internalization of ACKR3 and its delivery to the early endosomes. Of note, GRKs have been shown to be essential for efficient ligand-induced receptor internalization, as opposed to β-arrestins, which appear to be dispensable^31,32,34,35^. While constitutive ubiquitination of the receptor and interaction with RAMP3 were proposed to regulate the shuttling of ACKR3 to and from the plasma membrane^30,36,37^, the later stages of intracellular trafficking, including post-endocytic sorting, remain elusive. Within this context, whether ligand-induced and constitutive receptor cycling follow the same intracellular trafficking route and how this relates to the observed release of ACKR3-harboring vesicles has yet to be determined.

Hence, we investigated the importance of GRKs, β-arrestins, ubiquitination and RAMPs in both ligand-induced and basal EV release, using complementary approaches based on CRISPR-edited cell lines and ACKR3 mutants (**Fig. 6a**).

**Figure 6:**
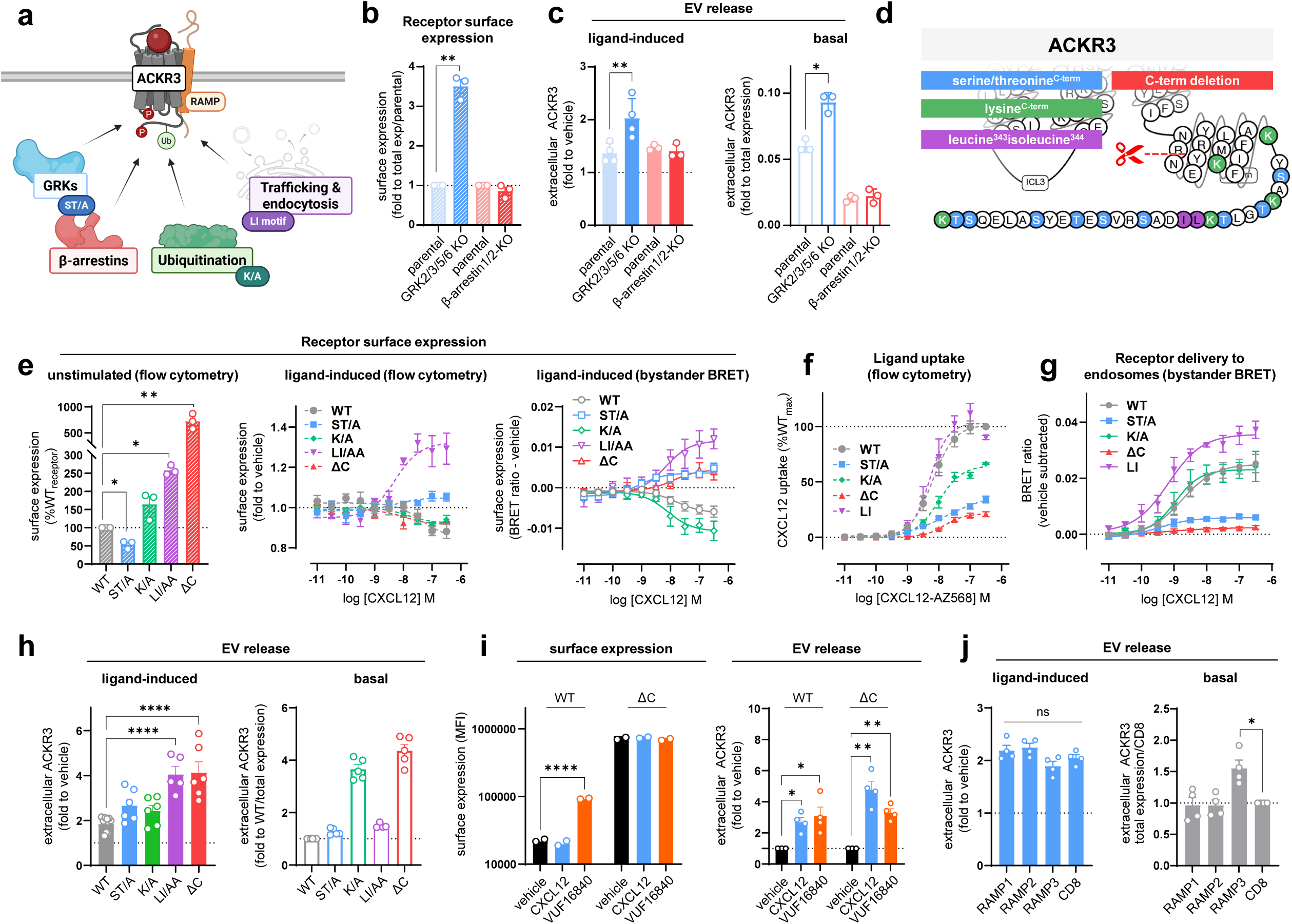
Ligand-induced and basal release is independent of GRKs and. β**-arrestins, and differentially affected by receptor C terminus motifs and RAMP interactions. (a)** Schematic representation of the key actors and residues/motifs involved in the trafficking of ACKR3. **(b)** Detection of ACKR3 at the plasma membrane of parental or CRISPR-edited HEK293 cell lines devoid of GRK2-3 and 5-6 (GRK2/3/5/6 KO) or β-arrestin 1 and 2 (β-arrestin 1/2 KO) monitored by flow cytometry using HiBiT-specific antibody. **(c)** Quantification of HiBiT-tagged WT ACKR3-harboring vesicles in the supernatant of parental, β-arrestin 1/2 KO or GRK2/3/5/6 KO cells after treatment with CXCL12 (*left*) or under basal conditions (*right*). **(d)** Schematic representation of the intracellular region of ACKR3, showing the C-tail (red delineation) and the localization of putative serine/threonine phosphorylation sites (blue), lysine ubiquitination sites (green) and the dileucine motif (LI) (purple). **(e)** Detection of WT ACKR3, or receptors with mutated serine and threonine (ST/A) or lysine (K/A) residues, the LI motif (LI/AA) or C-tail deletion (ΔC, Δ320-362) at the plasma membrane of transfected HEK293T cells in unstimulated conditions (*left*) or with increasing concentrations of CXCL12 (*middle*) monitored by flow cytometry using HiBiT-specific antibody and bystander NanoBRET (*right*). **(f)** Quantification of the uptake of fluorescently labelled CXCL12 (CXCL12-AZ568) by HEK293T cells expressing WT or mutated ACKR3 monitored by flow cytometry. **(g)** Endosome delivery of ACKR3 and mutants in response to increasing concentrations of CXCL12 assessed by bystander BRET. **(h)** Quantification of vesicles harboring WT or mutated/truncated HiBiT-ACKR3 in the supernatant after treatment with CXCL12 (*left*) or under basal conditions (*right*). **(i)** Detection of WT and ΔC mutant ACKR3 at the plasma membrane after 1 h incubation with CXCL12 and the inverse agonist VUF16840 (1 µM) monitored by flow cytometry using HiBiT-specific antibody (*left*). Quantification of vesicles harboring WT or ΔC ACKR3 in the supernatant after treatment with ligands using LgBiT-HiBiT complementation (*right*). **(j)** ACKR3-harboring vesicles in the supernatant of HEK293T cells expressing each RAMP after treatment with CXCL12 (*left*) or under basal conditions (*right*). CD8 was used as a negative control. CXCL12 was used at 100 nM unless otherwise stated. Results are presented as mean ± S.E.M of at least three independent experiments (n ≥ 3). *p < 0.05, **p < 0.01, ****p < 0.0001, by one sample t-test comparing a normalized value of 1 (b) or 100 (e), unpaired t-tests comparing parental vs KO (c), lognormal one-way ANOVA (i *right*), one-way ANOVA with Dunnet’s correction (h, j), two-way ANOVA with Šídák’s multiple comparisons test (i *left*). Panel (d) was created using GPCRdb.

CRISPR-edited HEK293 cells devoid of GRK2-3 and 5-6 (ΔGRK2/3/5/6 cells) or β-arrestin 1 and 2, revealed that the absence of GRKs resulted in a large increase of ACKR3 found in the supernatant under ligand-induced conditions, in line with the increased surface expression in these cells (**Fig. 6b–c)**. The removal of GRKs also increased the basal release, while the absence of β-arrestins did not influence the release of vesicles in either condition.

We then designed a series of ACKR3 mutants to clarify the impact of intracellular signal transducers, post-translational modifications and subcellular localization of the receptor on the release of EVs under ligand-induced and basal conditions. We generated mutations that prevent phosphorylation (ST/A) or ubiquitination (K/A) or truncated the C terminus of the receptor (ΔC). We also included a mutant of the “dileucine motif” (LL/I), present in the C terminus of ACKR3, and described to regulate internalization, post-endocytic trafficking and intracellular sorting of several GPCRs^65–67^ (**Fig. 6d**).

For ACKR3-ST/A, plasma membrane expression was reduced, alongside markedly impaired ligand-induced internalization (**Fig. 6e**) and diminished chemokine uptake capacity (**Fig. 6f–g, Supplementary Fig. 6**). Despite this, EV release remained comparable to WT under both ligand-induced and basal conditions (**Fig. 6h**). In line with the increased EV release observed in ΔGRK2/3/5/6 CRISPR-edited cells (**Fig. 6b–c**), these findings indicate that C-tail phosphorylation is not required for EV release and may instead restrain it by supporting receptor internalization and intracellular trafficking. In this context, ST/A likely recapitulates GRK deficiency by preferentially directing receptor toward EV release, though this effect is masked at the level of total EV release due to reduced basal receptor abundance at the plasma membrane.

Removing ubiquitination sites from the C terminus of ACKR3 (K/A) had the opposite effect on the surface expression of the receptor compared to the ST/A mutant, showing a slightly increased plasma membrane localization while retaining its ability to internalize and mediate ligand uptake (**Fig. 6e–g**). Interestingly, its ligand-induced EV release was not modified, whereas basal release exhibited a striking four-fold increase compared to the WT receptor (**Fig. 6h**), suggesting that ubiquitination of ACKR3 preferentially directs the receptor away from basal EV release pathways without affecting ligand-induced mechanisms.

In contrast, the ACKR3-LI/AA mutant exhibited even higher plasma membrane expression than WT or K/A mutant and maintained efficient ligand uptake (**Fig. 6e–g**). Upon stimulation, however, this mutant showed a surprising net increase in plasma membrane levels, alongside an enhanced ligand-induced vesicles release, with no detectable impact on basal release (**Fig. 6e, h**). This suggests that this motif is an important regulator of ACKR3 sorting, and that its modification, as opposed to the K/A mutation, preferentially steers the receptor toward the ligand-induced EV release pathways.

In line with the above findings, the deletion of the ACKR3 C terminus (ACKR3-ΔC) led to a marked increase in surface expression, accompanied by impaired ligand uptake, the latter being most likely due to the loss of phosphorylation sites (**Fig. 6e–g**). Despite this, ACKR3-ΔC exhibited a much stronger release into the extracellular compartment under both ligand-induced and basal conditions than the WT ACKR3 (**Fig. 6h**), recapitulating the combined effects observed with the LI/AA and K/A mutants respectively, both motifs being absent in this receptor variant. Notably, we observed that for this internalization-deficient mutant, the release of vesicles can also be induced by the inverse agonist VUF16840 (**Fig. 6i**), supporting the idea that a ligand-driven effect on ACKR3 conformation is needed to trigger the sorting of the receptor into EVs. The results also suggest that the increase in EV release following ACKR3 treatment with VUF16840 is two-fold, combining an increased presence of the receptor at the plasma membrane through inhibition of its basal cycling and a conformational change that is required to trigger vesicle release.

Finally, given that interaction with receptor activity-modifying proteins (RAMPs), notably RAMP3, was proposed to regulate the intracellular trafficking of ACKR3^36,37^, we wondered about the potential role of these single pass accessory proteins in EV release. We therefore assessed the impact of each individual RAMP on basal and ligand-induced release of ACKR3, relative to the negative control CD8. We found that while none of the RAMPs modulated the release of ACKR3 following ligand stimulation, the presence of RAMP3 significantly enhanced the basal release of the receptor suggesting that RAMP3 can also regulate/target ACKR3 toward basal EV release pathways (**Fig. 6j**).

These results further demonstrate that ligand-induced and basal ACKR3 release in EVs occur through distinct mechanisms involving different trafficking and regulatory pathways **(Fig. 7)**. Notably, neither process requires GRK-mediated phosphorylation or β-arrestin interaction, in contrast to ligand-induced receptor internalization and ligand uptake for which GRKs are essential. Instead, we confirmed that ligand-induced release is associated with receptor presence at the plasma membrane and influenced by later stages of receptor intracellular trafficking, whereas basal release appears to be regulated by ubiquitination and the trafficking chaperone RAMP3.

**Figure 7:**
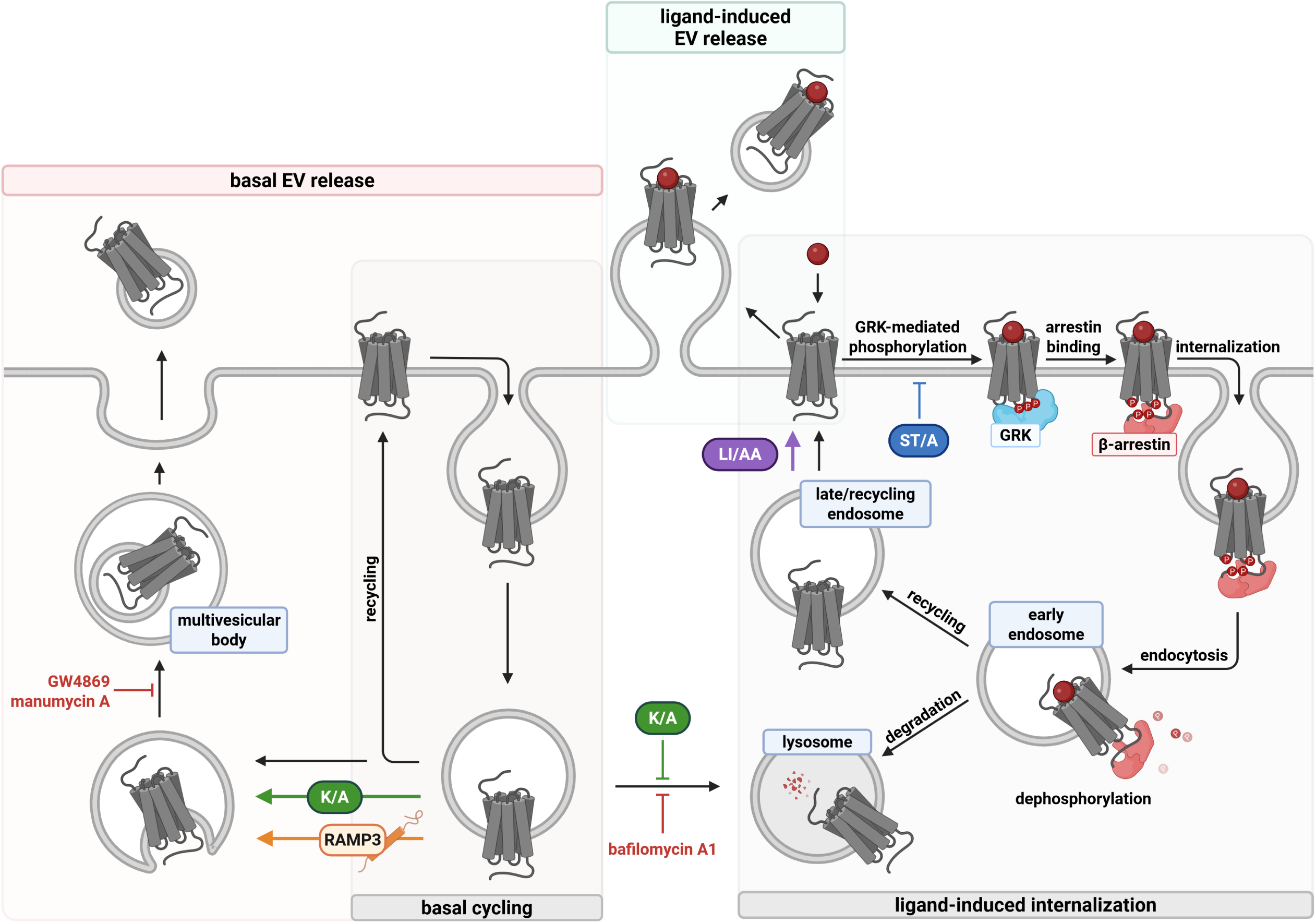
Model of ACKR3 trafficking supporting its scavenging function by internalization and release of extracellular vesicles. Schematic representation of two distinct putative ACKR3-EV release pathways. **Left**: In the absence of ligand, ACKR3 constitutively internalizes and either recycles back to the plasma membrane or is released on EVs. Blocking lysosome acidification by bafilomycin A1 or preventing ubiquitination of C-terminal lysine residues (K/A) enhances this release, while GW4869 and manumycin A reduce it. **Right**: Upon ligand binding, serine and threonine residues (ST) are phosphorylated by GRKs and β-arrestins are recruited. The receptor undergoes endocytosis, and either recycles back to the plasma membrane, is degraded in lysosomes or released on EVs. Mutation of the C-terminal LI motif (LI/AA) promotes receptor recycling to the plasma membrane and EV release.

Altogether, our findings suggest that EVs represent a previously underappreciated dimension of receptor trafficking and ligand regulation by ACKR3, with the potential to mediate both local and remote control of ligand availability.

## 3 Discussion

ACKRs form a subfamily of receptors unable to trigger G protein-dependent signaling cascades. Instead, they play a crucial regulatory role by controlling ligand availability^4–7^. ACKR3 is one of the best studied ACKRs and represents a valuable therapeutic target being involved in many physio- and pathological processes, including cancer and cardiovascular diseases^10,18,68^. Yet, many questions pertaining to the biology of ACKR3 remain to be elucidated, including the involvement of GRKs and β-arrestins in ACKR3 trafficking.

Over the years, research on GPCRs and chemokine receptors has benefitted from the ever-improving technologies and assays, allowing to continuously unveil new mechanisms and/or provide new interpretations for previously unexplained observations^69^. The present study illustrates this well with the use of recently developed techniques, namely HiBiT–LgBiT Nluc complementation and NanoBRET assays that led us to investigate an overlooked cellular compartment and dimension of ACKR3 trafficking biology.

We found that ACKR3 is released on EVs in the absence of ligand, and that this release is increased upon ligand stimulation. While neither basal nor ligand-induced release of ACKR3 on EVs requires GRKs or β-arrestins, the two processes are differentially regulated by determinants present in the receptor C terminus, such as ubiquitination sites or the dileucine motif or through interaction with accessory chaperone proteins known to regulate receptor trafficking, such as RAMP3. This suggests that ACKR3 release on EVs occurs via two independent trafficking routes and aligns well with recent reports that ACKR3, as well as other chemokine receptors, may exist as two populations with distinct trafficking fates **(Fig. 7)**^31,51^. The first population, tied to GRKs, undergoes internalization upon ligand binding, while the second cycles constitutively, independently of GRK or β-arrestin presence. ACKR3-harboring vesicles released under basal conditions likely originate from intracellular compartments of this continuously cycling pool, whereas ligand-induced EVs emerge upon receptor–ligand interactions at the plasma membrane **(Fig. 7)**. Additional studies will help conclusively elucidate the sorting pathways and the exact cellular machinery involved in the release of ACKR3 on EVs.

The observation that ACKR3 is released in EVs leads to two unexpected functional consequences. First, it becomes evident that upon binding, a proportion of ligand–ACKR3 complexes is directed to the extracellular space instead of being internalized. Their fate remains unclear but may include degradation or phagocytosis, transfer to neighboring cells^49,70–72^, or even long-distance cellular communication^73,74^. The potential ACKR3–CXCR4 heterodimers at the cell surface could also be impacted, as the two receptors are not co-released in EVs following ACKR3 stimulation. Second, the vesicles released under basal conditions harbor functional ACKR3 receptors capable of binding and, thereby, sequestering the ligand. These vesicles may trap ligands in the vicinity of the cells, reducing their local concentration and consequently limiting their availability for signaling receptors. Additionally, vesicles were also detected at greater distances from the releasing cells, suggesting possible remote scavenging capacities.

The contribution of this extracellular scavenging may also help understand previous data, especially those generated with internalization-deficient ACKR3 mutants and *in vivo* rescue experiments in mice and zebrafish^32,75^. Indeed, several independent studies showed that ACKR3-ST/A and ACKR3-ΔC fail to efficiently internalize and degrade CXCL12^29,31,76^. Consistently, mice expressing the phosphorylation-deficient ACKR3-ST/A mutant also display impaired CXCL12 internalization and scavenging. However, *Ackr3*^STA/STA^ variant did rescue the perinatal lethal *Ackr3*^-/-^ phenotype, although the specific mechanism involved was not elucidated^32^. Our study found that despite its reduced surface expression and impaired ability to internalize chemokines, ACKR3-ST/A is released on EVs and fully preserves its ability to bind CXCL12 in the extracellular environment. Hence, extracellular ligand sequestration on EVs could play a role in restoring the healthy phenotype.

This study reports EV release from multiple cell lines exogenously and endogenously expressing ACKR3, as well as primary cells. Future mechanistic and functional studies should seek to confirm the importance of the origin, nature and roles of ACKR3-harboring vesicles in additional relevant models (*e.g.* animal models of cancer^15,77^, multiple sclerosis, heart conditions^23^, pain^27^ or anxiety^78^). Importantly, the predominance of basal versus ligand-induced EV release is likely context-dependent and may vary across cell types and tissues, and therefore remains to be defined in vivo.

EVs have been associated with tumor progression and metastasis development in many cancer types^79–83^. Considering the described upregulation of ACKR3 in many tumors, it is tempting to speculate that the contribution of ACKR3-mediated extracellular scavenging may have an impact in cancer, notably through the CXCL12–CXCR4 axis^18,84,85^. Along this line, we found that different cancer-derived cell lines and CLL patient-derived cells released significant amount of ACKR3-harboring EVs. Additionally, bioluminescence signal in biofluids detected at remote sites from the initial orthotropic tumors in mice may be due to circulating ACKR3-EV pools.

The release of functional receptors on EVs may represent a more widespread mechanism than initially conceived, playing a pivotal role as part of the regulatory machinery. While a basal extracellular release was already proposed for other chemokine receptors, such as CCR5, CXCR4 and US28^44,46,70,72,86^, our study extends this concept to the dual atypical chemokine and opioid receptor ACKR3, for which both a ligand-induced and basal release could be demonstrated. Beyond ligand scavenging, receptor-harboring EV release may play a role in intercellular communication, receptor transfer/availability or ligand presentation at distal sites^40,44–49^. A recent report also demonstrated that stimulation of various receptors, including ACKR3, alters the miRNA content of EVs^87^. In support of this emerging paradigm, a recent study demonstrated that endothelial ACKR1 is secreted in EVs as a means of receptor downregulation following neutrophil-induced expression, further illustrating how receptors may exploit EV release to tune their surface availability^88^. EVs are also recognized as key players in the central nervous system^89–91^, including in the opioid system, where neuron-derived EVs have been shown to carry membrane-associated receptors, including KOR. Our study suggests that ACKR3-EVs may regulate opioid peptide levels such as BAM22 and may thus also participate more broadly in modulating opioid availability and pain-related signaling.

Finally, promoting receptor internalization over EV release may represent a novel approach for modulating ACKR3 function. Indeed, we found that while large endogenous ligands behave as ligands inducing both processes, small-molecule agonists such as VUF11207 and VUF25444 preferentially promote receptor endocytosis, whereas inverse agonists such as VUF16840 enhance EV release. These findings may have important implications for the development of future ACKR3-targeting compounds and are supported by recent structural studies suggesting that distinct ACKR3 modulators induce different receptor conformations involving differential engagement of the second extracellular loop^92,93^.

Overall, the findings of this study challenge the conventional understanding of ACKR3 biology, urging a more comprehensive exploration of its regulatory functions and their implications for intercellular communication. While ACKR3 is the only chemokine receptor to be released extracellularly upon activation, other receptors may be released in basal conditions and exhibit a range of functions, including but not limited to scavenging^40,44,46^. In a broader context, this study underscores the emerging paradigm of GPCR-harboring EVs and emphasizes the need to consider the often-overlooked extracellular compartment in tandem with the intracellular events^45^. To further our understanding, future research should delve deeper into the physiological repercussions of these EVs, the precise mechanisms governing their release, their composition in basal and ligand-induced contexts, as well as the varying nature of these particles across different cellular contexts. Additionally, investigations should explore the therapeutic potential of targeting GPCR-bearing EVs to either prevent/block their release or harness their potential to modulate ligand concentrations. These directions hold promise for new avenues in the field of GPCRs and specifically chemokine receptor biology in the context of therapeutic intervention^48^.

## 4 Material and methods

### 4.1 Peptides, chemokines, small molecules and antibodies

All fluorescently labelled and unlabelled chemokines were purchased from Protein Foundry except murine chemokine CXCL12, CXCL16 and XCL1 which were purchased from Peprotech. Peptides BAM22 and PAMP-12 were acquired from Phoenix Pharmaceuticals. VUF11207 was obtained from Tocris. VUF25444 and VUF16840 were synthesized by VU Amsterdam. Manumycin A was purchased from APExBio and GW4869 and bafilomycin A1 from Sigma-Aldrich. A list of antibodies used throughout the study can be found in **Supplementary Table 2**.

### 4.2 Cell lines and culture

HEK293T cells (Abcam, RRID:CVCL_0063), MDA-MB-468, HeLa wild-type and HeLa knock-in Nluc-ACKR3 cell lines^94^ were grown in Dulbecco’s modified Eagle medium (DMEM) supplemented with 10% fetal bovine serum (FBS, Sigma) and penicillin/streptomycin (100 units per ml and 100 μg per ml, Gibco). MCF-7 cells (ATCC, RRID:CVCL_0031) were grown in RPMI medium supplemented with 10% FBS and penicillin/streptomycin (100 units per ml and 100 μg per ml). HEK-HiBiT-ACKR3 cells were established by transfecting HEK293T cells with pIRES-puro vector encoding ACKR3 N-terminally fused with the HiBiT tag. Cells were maintained under puromycin selective pressure (5 μg per ml). β-arrestin and GRK knock-out cell lines were previously described^95,96^. U87 cells (ATCC) were grown in DMEM supplemented with 10% fetal bovine serum and penicillin/streptomycin (100 units per ml and 100 μg per ml).

### 4.3 Detection of receptor-harboring extracellular vesicle release by Nluc complementation

#### 4.3.1 Ligand-induced release of ACKR3-harboring vesicles

Receptor-positive EV release was measured using the Nano-Glo HiBiT Extracellular Detection System (Promega) according to the manufacturer’s protocol. HEK293T, HeLa, MCF-7 and U87-MG cells were seeded and 24 h later, when reaching ∼70% confluency, transfected with 1:10 plasmid encoding GPCRs N-terminally tagged with HiBiT, an 11-amino-acid tag that reconstitutes a functional Nanoluciferase upon high-affinity complementation with LgBiT. After further 24 h, cells were harvested, distributed in 96-well V-plates (1 × 10^5^ cells/well) and stimulated with the indicated ligand for 30 min at 37°C. The plates were then centrifuged (350 g × 5 min) and supernatants were transferred to white 96-well plates. Additional centrifugation steps were tested to establish an optimal protocol which confirmed lack of contamination by cell debris and efficient workflow (**Supplementary Fig. 1**). The Nano-Glo HiBiT extracellular reagent, containing the LgBiT protein (1:100) and Nanoluciferase extracellular substrate (1:50) was added and light emission upon the complementation between the two subunits was measured for 20 min using a GloMax Discover plate reader (Promega).

#### 4.3.2 Basal release of ACKR3-harboring vesicles

HEK293T cells were transfected as described in 4.3.1. After 24h cells were detached, washed, and resuspended in OptiMEM in V-bottom plates (1.5 × 10□ cells/well). Conditioned supernatants were collected at T_0_ = 5 min and 6 h, cleared by centrifugation (2,000 g × 5 min), transferred to white 96-well plates. The Nano-Glo HiBiT extracellular reagent was added and light emission upon the complementation between HiBiT and LgBiT was measured for 20 min using a GloMax Discover plate reader (Promega).

### 4.4 Detection of total receptor expression by Nluc complementation

Total cellular expression of HiBiT-tagged receptors was quantified using Nano-Glo HiBiT lytic detection system (Promega) according to manufacturer’s protocol. The HiBiT lytic reagent consisting of cell lysis buffer supplemented with Nanoluciferase substrate (1:50) and LgBiT protein (1:100) was added to cells and luminescence upon the complementation of the two subunits was measured for 20 min using a GloMax Discover plate reader (Promega).

### 4.5 Detection of surface receptors and markers by flow cytometry

Cell surface ACKR3 levels were determined by flow cytometry using ACKR3-specific mAb (12.5 μg per ml), clone 11G8 (R&D Systems, #MAB42273) or a matched isotype control (12.5 μg per ml) clone MG1-45 (BioLegend, #401402) or using anti-HA mAb (1 μg per ml) clone 16B12 or using anti-HiBiT mAb (2.5 μg per ml, Promega, #N7200) and allophycocyanin-conjugated F(ab’)2 fragment anti-mouse IgG (dilution 1:300, Jackson ImmunoResearch, #115-136-068). Dead cells were excluded using the Zombie Green fixable viability dye (BioLegend, dilution 1:3000, #423112). Fluorescence intensity was quantified on a Novocyte Quanteon flow cytometer (ACEA Biosciences) using NovoExpress 1.4.1 (ACEA Biosciences).

Surface marker detection on EVs was assessed following their immobilization on latex beads, as previously described^97^. Briefly, 5 µg of purified EVs were incubated with 10 µl of aldehyde/sulfate latex beads (#A37304, Thermo Fisher Scientific) for 15 min, after which PBS was added to a final volume of 1 ml and the mixture was incubated for 2 h on a rotating wheel at room temperature. Glycine was added to a final concentration of 100 mM and samples were incubated for an additional 30 min to block unoccupied binding sites. Samples were centrifuged (4,000 g × 3 min) and washed three times with PBS containing 0.5 % BSA. EV-coated beads were stained for 30 min at 4 °C with the following antibodies (1 µl/5 µg of EV): anti-ACKR3 (clone 8F11-M16, #331114), anti-CD63 (#353004), anti-CD81 (#349506), and anti-CD9 (#312105), all purchased from BioLegend. Beads were washed three times with PBS/0.5 % BSA and resuspended for acquisition on a Novocyte Quanteon flow cytometer using NovoExpress 1.4.1 (ACEA Biosciences).

For single-EV flow cytometry, 20µl of isolated EV samples were stained for 1 h at 4°C using the following antibodies (1 µl/5 µg of EV): anti-CD63 (#353011, BioLegend), anti-CD81 (#349506, BioLegend) and anti-ACKR3 (clone 8F11-M16, #331114, BioLegend). Stained EV samples were further diluted before acquisition to avoid swarm detection. MB488□positive events were used to define CD63, CD81 and ACKR3 gates. The ID7000 Spectral Cell Analyzer using ID7000 Software (Sony Biotechnology) was configured as previously described^98^ using the minimal flow rate and an acquisition time of 2 minutes per sample.

### 4.6 Uptake of fluorescently labelled chemokines monitored by flow cytometry

HEK293T cells were seeded in 10-cm culture dishes (6 × 10^6^ cells/dish) and 24 h later transfected with plasmids encoding wild-type or mutated ACKR3 N-terminally tagged with HiBiT. After further 24 h, cells were harvested and incubated for 30 min at 37°C with increasing concentrations of CXCL12-AZ568 (0.001 to 300 nM). Cells were then washed and treated with proteinase K for 2 h at 4°C to remove surface-bound chemokine, ensuring that only internalised signal was retained. Following additional washes, cell viability was assessed with a Zombie Green fixable viability dye (#423112, BioLegend, dilution 1:3000) and fluorescence intensity was measured on a Novocyte Quanteon flow cytometer using NovoExpress 1.4.1 (ACEA Biosciences).

### 4.7 Purification of receptor-harboring extracellular vesicles by size exclusion chromatography

Size exclusion chromatography (SEC) columns were made by stacking Sepharose CL/2B (GE Healthcare, #17-0140-01) in PBS up to a 10 ml bed volume. Supernatants were harvested from a single confluent 10-cm culture dish per condition or, for dot blot analysis with cell lines MCF7, HeLa, MDA-MB-468, from a confluent T175 flask. Following centrifugation steps to remove dead cells and cell debris (2 × 5 min centrifugation at 500 g and 2 × 10 min at 2,000 g), culture supernatant was loaded on the SEC column and allowed to enter by gravity flow. 0.5-ml fractions were collected immediately, and the EV-enriched and protein-enriched fractions were used for bioluminescence detection, NanoBRET chemokine binding with fluorescently labeled ligand or dot blot detection. For CLL patient samples, 10 × 10^6^ PBMC were isolated as previously described^99^ and seeded with 7.5ug/mL of ODN-2006 (#tlrl-2006-1, Invivogen). The supernatant was harvested after 24h and centrifuged 4 times (2 × 5 min centrifugation at 500 g and 2 × 10 min at 2,000 g), before freezing at -80°C.

### 4.8 Purification of receptor-harboring extracellular vesicles by differential ultracentrifugation

For basal EV release, stably expressing HiBiT-ACKR3 HEK293T cells were grown to 60% confluency in DMEM (four T175 flasks) then washed with PBS. After 48 h incubation in Opti-MEM, supernatants were collected. EVs were isolated following sequential centrifugation steps adapted from Gargiulo *et al*.^100^ (500 g × 5 min, 500 g × 20 min, 2,000 g × 40 min and 16,000 g × 60 min) to remove cell debris and larger vesicles. For ligand-induced release, cells were grown to 90% confluency in 28 10-cm culture dishes per condition (2 × 10^7^ cells/dish), washed the next day, and incubated in Opti-MEM in the presence or absence of BAM22 (1 µM). Supernatants were harvested after 6 h and processed as below.

EVs were isolated by ultracentrifugation using an Optima MAX-XP ultracentrifuge (Beckman Coulter). EVs were pelleted using an MLA-55 fixed-angle rotor at 110,000 g for 75 min at 4°C. The pellet was resuspended in 1 ml of PBS and subjected to lipid dye staining using BioTracker MemBright 488 Live Cell Dye (#SCT083, Sigma-Aldrich) for 1 h. To remove free dye and further purify the EV preparation, a top density cushion consisting of 17% OptiPrep (#1893, Progen) solution was used. Ultracentrifugation was performed using an MLS-50 swinging-bucket rotor at 110,000 g for 75 min with minimal acceleration and braking to avoid disrupting the EV band. EVs were collected and washed with PBS before a final ultracentrifugation pelleting step using the same parameters as the initial step. EVs were resuspended in PBS and stored at −80°C for further characterization.

### 4.9 Western blot, dot blot and protein quantification

EV preparations isolated through ultracentrifugation were mixed with 4X Laemmli buffer and loaded on 13% SDS–PAGE gels. Proteins were transferred to nitrocellulose membranes by wet transfer at 100V for 75 min. For dot blot, 100–200 µl of EV fractions were applied to nitrocellulose membranes using a Bio□Dot® apparatus (Bio□Rad). For HiBiT detection, the Nano□Glo® HiBiT Blotting System (#N2410, Promega) was used according to the manufacturer’s instructions. Ponceau S staining was used to assess total protein loading. Membranes were blocked for 1 h at room temperature in 5% milk in TBS□T under gentle agitation and incubated overnight at 4 °C with primary antibodies under agitation: anti-ACKR3 (#MAB42273, R&D Systems), anti□CD63 (#556019, BD Biosciences), anti□CD81 (#sc-7637, Santa Cruz) and purity controls anti□calnexin (ER-bound, #2433, Cell Signaling), and anti□PHB1 (marker of mitochondria-bound, #60092-1-Ig, Proteintech). After three washes in TBS□T, membranes were incubated for 1 h at room temperature with HRP□conjugated secondary antibodies (anti□mouse IgG #115-035-008 and anti□rabbit IgG #111-035-003, Jackson ImmunoResearch) under agitation. After washing, proteins were detected using SuperSignal™ West Femto Maximum Sensitivity Substrate (#34096, Thermo Scientific). Images for dot blot and HiBiT detection signals were acquired using an ImageQuant™ LAS 4000 system, and western blot signals were detected on films (#SC-201696, Santa Cruz). Band and spot intensities were quantified in ImageJ by measuring after background subtraction the pixel intensity.

### 4.10 Microfluidic nanoscale fluorescence-activated analysis

Fluorescent EV sample content was measured using a single-EV high-sensitivity nanoscale fluorescence-activated object sorting technology (HEKAT, France, www.hekat.com). PBS and PBS with working concentrations of ACKR3 antibody (clone 8F11-M16, #331114, BioLegend) were used as negative controls. Analysis volume (15 µl) was loaded on a single-use quantification microchip (model: 12 channels, flow rate; 5 µl/h, lasers: 633 nm) and transferred in the analyzer. After the automatic noise thresholding, fluorescent signals were recorded for 2 min. Measures were made in triplicates.

### 4.11 Extracellular vesicles size distribution and particle concentration

EV size distribution and particle concentration were measured on a Zetasizer Advance Ultra using ZS Explorer (Malvern Panalytical). Samples were collected and isolated by sequential centrifugation (500 g × 5 min, 2,000 g × 10 min) followed or not by SEC fractionation, or by ultracentrifugation (described in 4.8). Particle size was determined by multi-angle dynamic light scattering (MADLS), and particle concentration was estimated using the instrument’s particle concentration measurement mode.

### 4.12 Detection of receptor intracellular trafficking by NanoBRET

Ligand-induced receptor trafficking was assessed by bystander NanoBRET based on protocols described in greater detail in previous publications^101,102^. In both assays, HEK293T cells (6 × 10^6^) were seeded in 10-cm dishes and 24 h later co-transfected with plasmids encoding ACKR3 C-terminally tagged with Nanoluciferase (BRET donor) and an mNeonGreen-fused subcellular marker (BRET acceptor). To monitor receptor disappearance from the plasma membrane, NeonGreen was fused to the first 11 residues of human Lyn kinase, which anchors it to the inner leaflet of the plasma. For early endosome delivery, NeonGreen was fused to the PI3P-binding FYVE domain of endofin, which localises constitutively to early endosomes.

After further 24 h, cells were harvested, distributed into white 96-well plates (1 × 10^5^ cells/well), and stimulated with the indicated ligand for 1 h before addition of coelenterazine H (4 ng/ml). Donor (450/8 BP) and acceptor (530 LP) emissions were measured on a GloMax Discover plate reader (Promega) and results expressed as BRET ratio (acceptor/donor emission).

### 4.13 Detection of β-arrestin recruitment by NanoBiT

Ligand-induced β-arrestin recruitment was monitored by Nanoluciferase complementation assay (NanoBiT, Promega). Briefly, 6 × 10^6^ HEK293T cells were seeded in 10-cm culture dishes and 24 h later cotransfected with vectors encoding GPCRs C-terminally tagged with SmBiT and human β-arrestin 1 N-terminally fused to LgBiT. 24 h after transfection, cells were harvested, incubated 25 min at 37 °C with Nano-Glo Live Cell substrate diluted 200-fold and distributed into white 96-well plates (1 × 10^5^ cells per well). Ligands were then added at indicated concentrations and β-arrestin recruitment to the receptor was evaluated by measuring luminescence emitted upon Nanoluciferase complementation for 20 min with a GloMax Discover plate reader (Promega).

### 4.14 Detection of receptor-harboring extracellular vesicle release by NanoBRET

Formation or release of ACKR3–CXCL12 complexes or capture of free CXCL12 by ACKR3-harboring extracellular vesicles, were monitored by NanoBRET using AZ568-labelled CXCL12 (CXCL12-AZ568) as the acceptor and ACKR3 N-terminally fused to Nanoluciferase (Nluc) or the HiBiT tag complemented by LgBiT as the donor. Nluc substrate or HiBiT extracellular reagent (Nluc substrate + LgBiT protein) was added immediately prior to reading. Donor (450/8 BP) and acceptor (600 LP) emissions were measured on a GloMax Discover plate reader (Promega). For all experiments, 6 × 10^6^ HEK293T cells were seeded in 10-cm dishes and 24 h later transfected with plasmids encoding ACKR3 N-terminally tagged with HiBiT or Nluc, unless otherwise stated. Experiments were performed 24 h after transfection.

#### CXCL12-induced release of ACKR3-harboring vesicles

Transfected cells were harvested, washed, and distributed in 96-well V-bottom plates (1 × 10^5^ cells/well). After incubation with CXCL12-AZ568 (0.01 to 100 nM, 37°C, 30 min), plates were centrifuged (350 g × 5 min) and supernatants transferred to white 96-well plates for BRET measurement.

#### Binding of CXCL12 to ACKR3-harboring vesicles

Conditioned supernatants were collected 24 h after transfection, centrifuged (3,000 g × 10 min), and transferred to white 96-well plates. CXCL12-AZ568 (0.01 to 100 nM) was added and samples were incubated on ice for 2 h prior to BRET measurement.

#### Binding of CXCL12 to ACKR3-harboring vesicles prior to SEC

HEK293T or HeLa cells transfected with plasmids encoding Nluc- or HiBiT-tagged ACKR3 or knock-in Nluc-ACKR3 HeLa cells were cultured to 90% confluency in a 10-cm culture dish, at which point they were detached with trypsin, washed with PBS, resuspended in 2 ml Opti-MEM and incubated for 30 min at 37°C. Conditioned medium was subjected to sequential centrifugation steps (500 g × 5 min and 2,000 g × 10 min), supplemented with 200 nM CXCL12-AZ568, and rotated at room temperature for 1 h before SEC fractionation (as described in Section 4.6). BRET was measured in each fraction following substrate addition.

#### Binding of chemokine to receptor-harboring vesicles in pooled SEC fractions

Following SEC fractionation of conditioned supernatants from HEK293T cells transfected with plasmids encoding Nluc- or HiBiT-tagged receptors, EV-enriched fractions (fractions 7–11) and soluble protein fractions (fractions 16–20) were pooled separately. CXCL12-AZ568 (0.01 to 100 nM) was added to each pool prior to BRET measurement.

### 4.15 Ligand scavenging assay

To collect extracellular vesicles from the supernatant, 3 × 10^6^ HEK293T cells were seeded in ten 10-cm culture dishes and 24 h later transfected with plasmids encoding ACKR3 N-terminally tagged with HiBiT or an empty plasmid. 72 h after transfection, the supernatant was harvested, centrifuged (3 000 g × 10 min) and concentrated using Centricon® Plus-70 (100 kDa cutoff) (Millipore, Merck) according to manufacturer’s instructions.

For the scavenging assay, 6 × 10^6^ HEK293T cells were seeded in 10-cm culture dishes and 24 h later cotransfected with plasmids encoding CXCR4 C-terminally tagged with SmBiT and β-arrestin 1 N-terminally tagged with LgBiT (according to *4.13*) or with plasmids encoding untagged CXCR4 or MOR, Gα_i_ fused to the Nanoluciferase in the unstructured region after the second helix and Gβγ fused to cpVenus^103^.

The concentrated supernatant of cells expressing ACKR3 (ACKR3 EVs) or non-transfected cells (vehicle EVs) was incubated with increasing concentrations of CXCL12 and BAM22, ranging from 0.3 to 100 nM and 10 nM to 3 μM, respectively, for 3 h at 37°C with regular mixing steps.

For the β-arrestin assay, the cells were detached and pre-incubated with Nano-Glo Live Cell substrate diluted 200-fold for 25 min at 37 °C before adding the concentrated supernatant. Ligand-induced β-arrestin recruitment was measured with a GloMax Discover plate reader (Promega) for 20 min.

For the G protein dissociation assay, the cells were detached and plated in a 96-well plate. The concentrated supernatant was added before distributing Nano-Glo Live Cell substrate diluted 200-fold. Ligand-induced G protein dissociation was monitored with a GloMax Discover plate reader (Promega) for 30 min using a 450/8 BP filter (Nanoluciferase donor) and a 530 LP filter (cpVenus acceptor).

### 4.16 Super-resolution microscopy

#### 4.16.1 Antibody preparation

Antibodies against ACKR3 (#331102, Biolegend) were labelled with Alexa Fluor 647 using an Alexa Fluor antibody labeling kit following manufacturer’s instructions (#A88068, Thermo Fisher Scientific). In short, antibodies were re-equilibrated in appropriate buffer using a Zeba Dye and biotin removal spin columns (#A44296S, Thermo Fisher Scientific) following manufacturer’s instructions. Next, when applicable, antibodies were diluted to a concentration of 1 mg/mL and added a 1/10 volume of 1 M sodium bicarbonate buffer. The protein solution (100 μL) was then added to the reactive dye vial, inverted a few times and allowed to incubate for 15 minutes at RT. Meanwhile, spin columns were prepared for the purification of labelled proteins. The columns were centrifuged (1,000 g × 2 min) to allow for flowthrough removal. The column was then re-equilibrated by adding 500 μL of PBS Exchange Buffer and centrifuged (1,000 g × 2 min). The flowthrough was discarded and the column was placed into a fresh snap-cap collection tube. Following the incubation, the reaction mixture was carefully pipetted onto the center of the column and centrifuged (1,000 g × 2 min). The final eluted conjugated antibodies were stored based on the original storage conditions.

#### 4.16.2 Image acquisition

Super-resolution images of EVs were acquired using direct stochastic optical reconstruction microscopy (dSTORM; Nano-imager S, Oxford Nanoimaging (ONI)), equipped with a 100X, 1.4NA oil immersion objective. The EV Profiler Kit 2 (ONI) was used per manufacturer’s instructions. Surface capture was performed using the manufacturer’s capture reagent (calcium based). Image acquisition was performed in total reflection fluorescence (TIRF) mode with a 45–50° angle at 32°C. Laser excitation was performed with laser powers set to 30% and 50% for 647 nm and 561 nm channels, respectively. Calibration was carried out before imaging using TeTraSpek microsphere (#T7279, Thermo Fisher Scientific) to ensure channel alignment within a 10-nm standard deviation. 2,000 frames were acquired (1,000 for each channel) with a 30-ms exposure using AutoEV in the collaborative discovery (CODI) software. Localization images were drift-corrected, filtered by photon count and localization precision and subsequently clustered and quantified to obtain population statistics and relative fluorophore intensity.

### 4.17 Human subjects and ethics

All experiments involving human samples were conducted in accordance with the Declaration of Helsinki and approved by the relevant institutional ethics committees, including the Luxembourg Comité National d’Ethique et de Recherche (CNER N°202407/03) and the Ethik-Kommission an der Medizinischen Fakultät der Eberhard-Karls-Universität sowie am Universitätsklinikum Tübingen (approval 597/2023BO2). Written informed consent was obtained from all participants prior to sample collection. CLL patients were randomly picked from the ongoing TSI-2.0 study, with no specific recruitment criteria. All study procedures complied with established good clinical practice guidelines.

### 4.18 Detection of EV released by platelets

The preparation of isolated human platelets for the ACKR3 surface expression was executed as described previously^104^.

For ACKR3 surface expression, isolated platelets were adjusted to a count of 1 × 10^6^ per sample in Dulbecco’s phosphate buffered saline (0.5 mM MgCl_2_, 0.9 mM CaCl_2_). The sample was stained with an ACKR3-specific antibody (8 µg/ml, clone 8F11-M16, #331114, BioLegend) or an isotype control for 30 min at room temperature. Cells were fixed with 300 µl 0.5 % paraformaldehyde and analyzed on FACSLyric™ using BDSuite™ (BD Bioscience) flow cytometer.

For the EV measurements, citrate-anticoagulated blood was centrifuged (209 g × 20 min without brakes) to obtain platelet-rich plasma (PRP). The PRP was harvested and the cell count was determined using a SYSMEX cell counter (Sysmex Cooperation). The remaining sample was centrifuged (3,300 g × 10 min) to obtain platelet poor plasma (PPP). The platelet count was adjusted to 200,000 platelets per µl with PPP.

PRP were adjusted to 200,000 platelets per µl with PPP and 150 µl of the suspension was incubated in a glass cuvette for 10 min at 37°C and 1,000 rpm stirring with or without chemokine. 10 µl of the sample was stained with anti-ACKR3 antibody in a 50 µl assay for 30 min at RT. Cells were then fixed with 300 µl 0.5 % paraformaldehyde.

For the antibody competition assay, samples were preincubated 15 min with the following antibodies: anti-ACKR3 (8 µg/ml, clone 8F11-M16, #331102, BioLegend), anti-CXCR4 (8 µg/ml, clone QA18A64, #304503, BioLegend), and their respective isotype controls. Then, 250 nM CXCL12 was added and incubated for 10 min at 37°C and 1,000 rpm stirring. Again, 10 µl of sample were diluted with 300 µl 0.5 % paraformaldehyde and analyzed by flow cytometery. The EV measurements were performed with a FACSLyric™ (BD Bioscience, San Jose, CA, USA) flow cytometer. A suspension of 1 µm beads was measured to obtain a 1 µm cut off. The EV percentage represents the fraction of events smaller than 1 µm compared with the total event count. The raw data of all flow cytometer measurements were analyzed using FlowJo software (version 10.10.0 FlowJo CCL).

### 4.19 Animal experiments

All experiments involving laboratory animals were conducted in a pathogen-free animal facility with the approval of the Luxembourg Ministry for Agriculture (#TSI-2020-01). Mice were treated in accordance with the European Union guidelines. NSG mice (MGI: 3577020, RRID:IMSR_JAX:005557) were obtained from Jackson Laboratories. 1 × 10^6^ WT or Nluc-ACKR3 HeLa cells were resuspended in 100 µl of a 1:1 DMEM/Matrigel (Corning™ 354234) mixture and injected subcutaneously in the rear flank of the mice. After sufficient tumor development, mice were euthanized by cervical dislocation. Peripheral blood was collected before euthanasia and placed in EDTA-containing tubes. Plasmas were prepared by centrifugation at 1500 g, 10 min at 4°C and then stored at -80 °C until further analysis. Tumors were weighed using a precision scale and were placed on nylon mesh filter taped on top of a 50 mL conical tubes and centrifuged at 4 °C, 10 min at 100 g. The tumor interstitial fluid was collected and used for further experiments. The dissociation of the spleen was performed using 100 µm cell strainer. The resuspension volume ratio was 1 ml of PBS for 10 mg of spleen cell suspension. The cell suspension was centrifuged for 5 min at 400 g. The collected supernatant was centrifuged at 1,000 g for 5 min. The final supernatant was filtered on 0.22 µm filter. The release of ACKR3-harboring vesicles from the tumors was determined by quantifying the Nluc luminescence in the plasma, spleen supernatant and tumor interstitial fluid using GloMax Discover plate reader (Promega).

### 4.20 Statistics

Concentration–response curves were fitted to the four-parameter Hill equation using an iterative, least-squares method (GraphPad Prism version 10.6.1). All curves were fitted to data points generated from the mean of at least three independent experiments.

All statistical tests were performed with GraphPad Prism 10.6.1. P-values are indicated as follows: *p < 0.05, **p < 0.01, ***p < 0.001, **** p<0.0001.

### 4.21 Figure preparation

Schematic figures were created with BioRender (BioRender.com) and exported under a paid publication licence.

## Data availability

The datasets generated during and/or analyzed in the current study are available from the corresponding author on reasonable request.

## Acknowledgements

This study was supported by the Luxembourg Institute of Health (LIH) through the NanoLux Platform, the Cancer Foundation Luxembourg, Luxembourg National Research Fund (INTER/FNRS CXCL12 20/15084569, CORE IMPACTT C23/BM/18068832, CORE EVIL C20/BM/14592342, ACKROS C25/BM/19563724, INTER/AUDACE/25/19352183, PRIDE 19/14254520/i2TRON, and 21/16763386/CANBIO2, 16749720/NextImmune2 and AFR Opiokine 14616593), F.R.S.-FNRS-Télévie (grants 7.4593.19, 7.4529.19, 7.8504.20, 7.4502.21 and 7.8508.22). CC is supported by The Netherlands Organisation for Health Research and Development (ZonMw) [ZonMw Veni; 09150162010212, and ZonMw Off-Road; 04510012110012]. LN is supported by The Netherlands Health Research Organisation (ZonMw) grant (ZonMw Open Competition; 09120012110079). AK is supported by The German Research Foundation (DFG) – Project number 552766849. This project was supported by the German Research Foundation (DFG) – Project number 335549539 / GRK 2381 and 374031971 / TRR 240 to MG. MB was supported by a discovery grant (# RGPIN-2019-05556) from the Natural Sciences and Engineering Research Council of Canada. CBP, SJH, CH, RL, MJS, MS, AC are part of the Marie Skłodowska-Curie Innovative Training Network ONCORNET2.0 “ONCOgenic Receptor Network of Excellence and Training” (MSCA-ITN-2020-ETN, Program under Grant Agreement 860229). The authors would like to thank Jean-Marc Plesseria, Nadia Beaupain, Sandrine Pierson, and Dr. Antonio Cosma, Mario Gomez, Maria Konstantinou from the Luxembourg National Cytometry Platform, Anaïs Oudin and Coralie Pulido from the Animal Facility staff for technical and experimental support, the CORE Extracellular Vesicles facility (Ghent University, Belgium), as well as Wilfrid Mazier and Selma Cornillot-Clément at Hekat for providing early access to the nanoscale fluorescence-activated object sorting technology prior to commercial release and for sample processing.

## Authors contributions

CBP, MS and AC designed the study and wrote the manuscript. LR, EM and JP contributed to the design and supervision of the study and to the writing of the manuscript. CBP, LR, MM, CC, MC, A-KR, LDN, AAB, CP, VK and EC performed the experiments. CBP, LR, MM, EM, JP, MJS, MS and AC analyzed and interpreted the data. CBP, LR, MM, JD and CH generated molecular tools for cellular assays. RL, SJH, MB, SAL, JD, CH, MPG and AH contributed essential material. AC, MS, EM and JP supervised the overall study. All authors reviewed and approved the manuscript.

