## Supplementary Data and Tables for "Chemokine and opioid peptide scavenging through constitutive and ligand-induced release of ACKR3-bearing extracellular vesicles"

### – Supplementary information –

Christie B. Palmer[1,3,‡], Lucie Rospape[2,3,‡], Max Meyrath[1], Caitrín Crudden[4], Manuel Counson[1], Anne-Katrin Rohlfing[5], Lotte Di Niro[4], Ana Alonso Bartolome[1,3], Cláudio Pinheiro[6,7], Vanessa Klapp[2,3], Ester Cassano[1,3], Stéphane A. Laporte[8], Stephen J. Hill[9], Julia Drube[10], Carsten Hoffmann[10], Rob Leurs[4], Michel Bouvier[11], Meinrad P. Gawaz[5], An Hendrix[6,7], Etienne Moussay[2], Martine J. Smit[4], Jérôme Paggetti[2†], Martyna Szpakowska[1†], Andy Chevigné[1,†]

[1] Department of Infection and Immunity, Luxembourg Institute of Health (LIH), Esch-sur-Alzette, Luxembourg
[2] Tumor Stroma Interactions, Department of Cancer Research, Luxembourg Institute of Health (LIH), Luxembourg
[3] Faculty of Science, Technology and Medicine, University of Luxembourg, Esch-sur-Alzette, Luxembourg
[4] Department of Chemistry and Pharmaceutical Sciences, Division of Medicinal Chemistry, Amsterdam Institute for Molecules, Medicines and Systems (AIMMS), Vrije Universiteit, Amsterdam, The Netherlands
[5] Department of Cardiology and Angiology, University Hospital Tübingen, University Tübingen, Tübingen, Germany.
[6] Laboratory of Experimental Cancer Research, Department of Human Structure and Repair, Ghent University, Ghent, Belgium
[7] Cancer Research Institute Ghent (CRIG), Ghent, Belgium
[8] Department of Medicine, Research Institute of the McGill University Health Center, and Department of Pharmacology and Therapeutics, McGill University, Montréal, Canada
[9] Cell Signalling and Pharmacology Research Group, Division of Physiology, Pharmacology and Neuroscience, School of Life Sciences, University of Nottingham, Nottingham, UK
[10] Institute for Molecular Cell Biology, CMB-Center for Molecular Biomedicine, University Hospital Jena, Friedrich Schiller University Jena, Jena, Germany
[11] Institute for Research in Immunology and Cancer (IRIC), Department of Biochemistry and Molecular Medicine, Université de Montréal, Montréal, Canada
‡: these authors contributed equally to this work
†: these authors contributed equally to this work

**Content:**

Supplementary Figures 1–6
Supplementary Tables 1–2

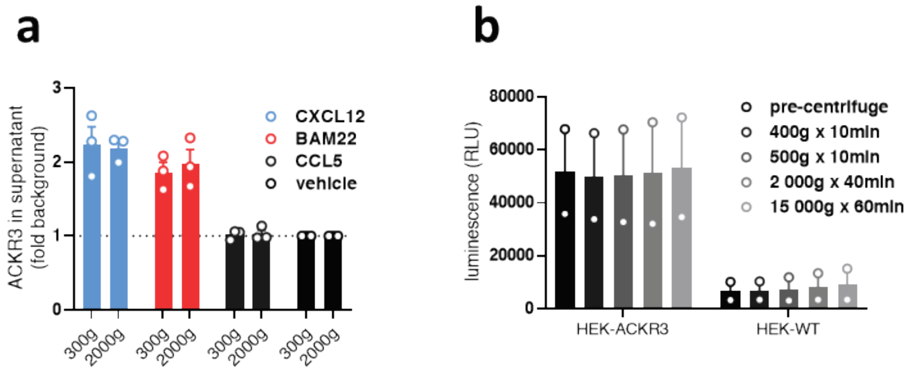


**Supplementary Figure 1**: **(a)** Detection of HiBiT-tagged ACKR3 in the supernatant of HEK293T cells stimulated with CXCL12 (100 nM), BAM22 (1 µM) or CCL5 (100 nM) isolated with a 5-min centrifugation at 300 g or 2,000 g. **(b)** Luminescence signal of the supernatant of unstimulated HEK293T cells expressing HiBiT-ACKR3 following various centrifugation steps. Results are presented as mean ± S.E.M of at least two independent experiments (n ≥ 2).

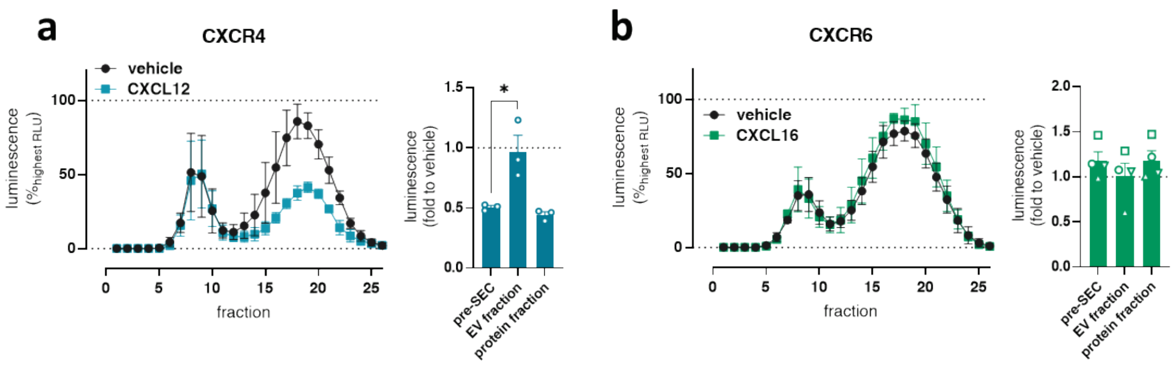


**Supplementary Figure 2**: SEC fractions from supernatants isolated from HEK293T cells transiently expressing N-terminally HiBiT-tagged CXCR4 (**a**) or CXCR6 (**b**) and stimulated or not with CXCL12 (100 nM) or CXCL16 (100 nM), respectively (*left*). Luminescence signal following stimulation in either total supernatant (pre-SEC) or pooled SEC fractions corresponding to EV-enriched or soluble protein-enriched fractions (*right*). Results are presented as mean ± S.E.M of at least three independent experiments (n ≥ 3). *p < 0.05 by one-way ANOVA with Dunnett's multiple comparison test.

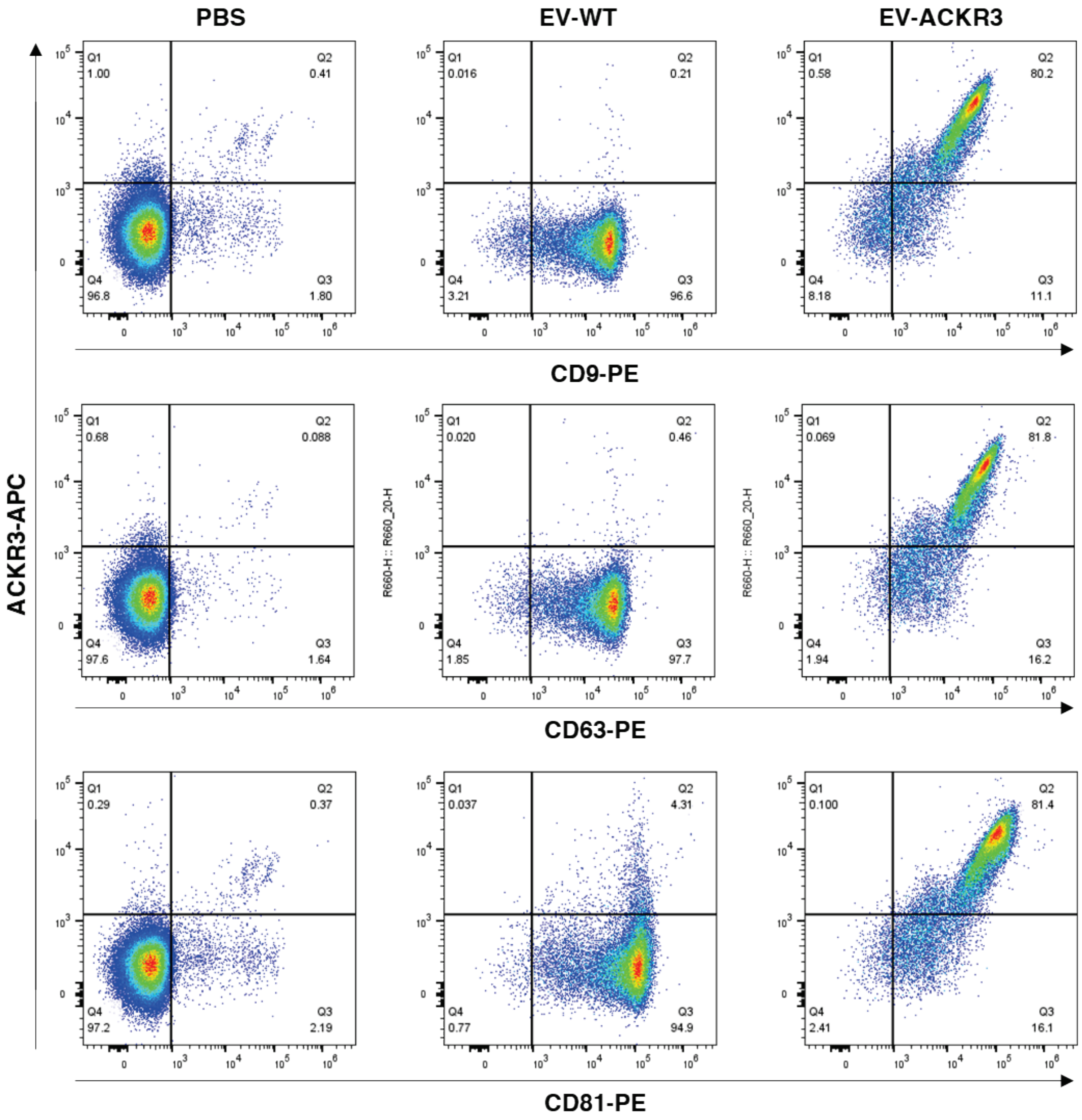


**Supplementary Figure 3**: Representative flow cytometry dot plots of aldehyde/sulfate latex microbeads coated with EVs from EV-WT and EV-ACKR3 preparations obtained under basal conditions isolated using ultracentrifugation, showing ACKR3 signal (y-axis, detected with 8F11-M16 antibody) against CD9, CD63 and CD81 signal (x-axis). PBS-coated beads incubated with antibody were used as a control to set the gates.

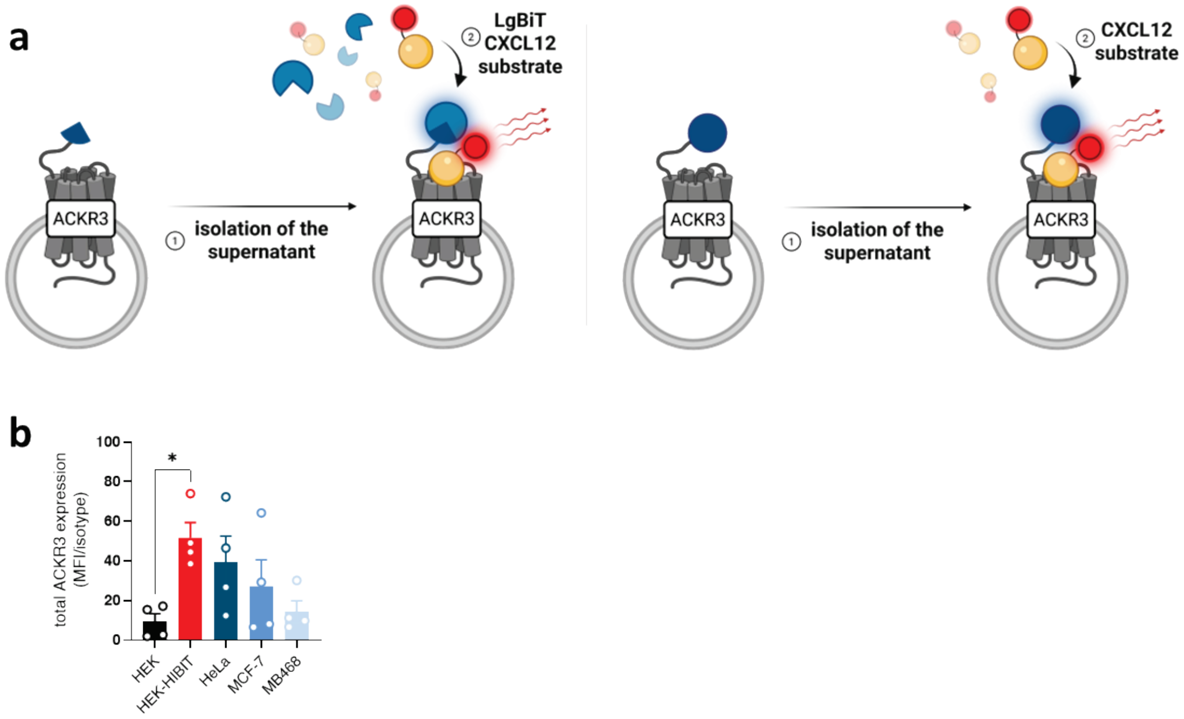


**Supplementary Figure 4**: (**a**) Schematic representation of the NanoBRET assay used to measure chemokine binding to ACKR3 on EVs released under basal conditions. Measurements were performed using either the full-length Nluc N-terminally tagged to ACKR3 (main Fig. 4a–b, right) or Nluc complementation via HiBiT-LgBiT association (main Fig. 4b, left and c): (1) isolation of supernatants from HEK293T cells expressing HiBiT- or Nluc-ACKR3, (2) addition of LgBiT (where applicable), CXCL12-AZ568 and substrate, followed by NanoBRET measurement. **(b)** Total ACKR3 expression monitored by flow cytometry in WT HEK293T or HEK293T transiently expressing HiBiT-ACKR3 and three cell lines (HeLa, MCF-7, MDA-MB-468) endogenously expressing ACKR3. Results are presented as mean ± S.E.M of four independent experiments (n = 4). *$p < 0.05$ by one-way ANOVA with Dunnett's multiple comparison test.

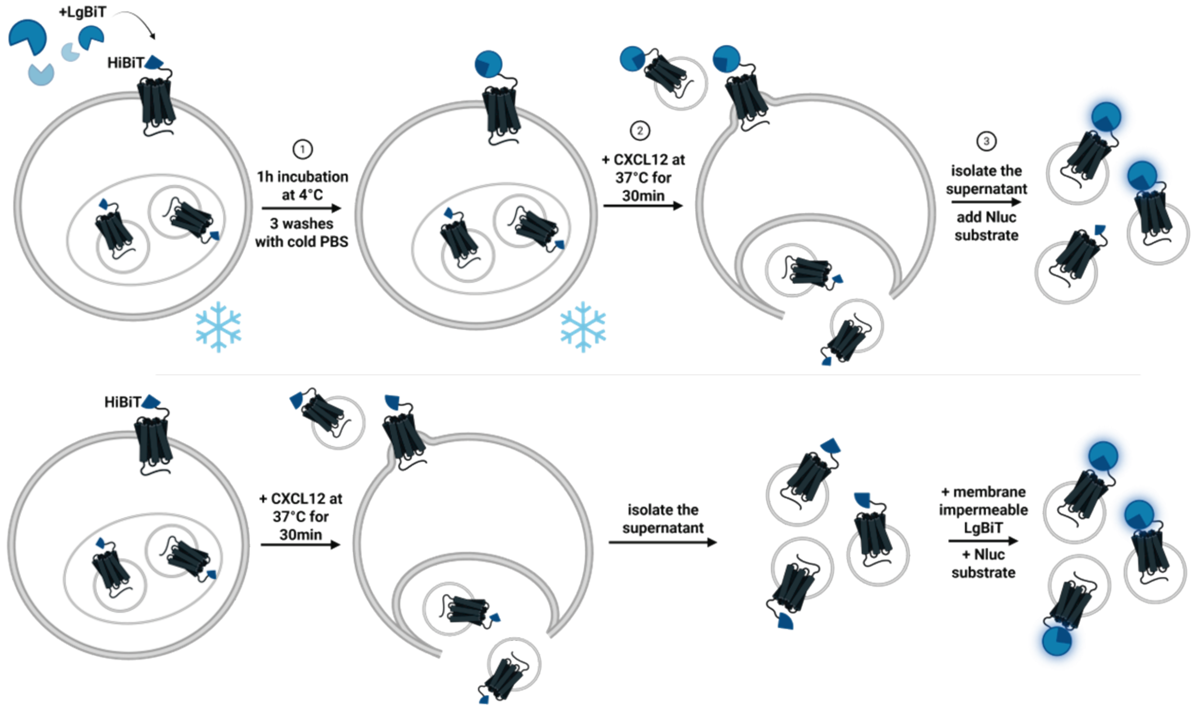


**Supplementary Figure 5**: Schematic representation of the assay demonstrating the surface origin of EVs released in response to ligand (*top*) (1) Incubation of cells expressing HiBiT-tagged ACKR3 with membrane-impermeable LgBiT for 1 h at 4°C to allow Nluc complementation in the absence of receptor trafficking, (2) ligand stimulation following extensive washes to remove unbound LgBiT, (3) supernatant isolation and quantification of luminescence upon addition of Nluc substrate. *(bottom)* Schematic representation of the conventional HiBiT assay monitoring the release of ACKR3-harboring EVs.

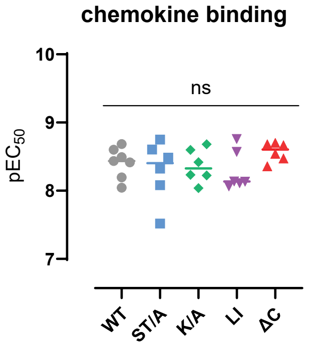


**Supplementary Figure 6**: CXCL12-AZ568 binding affinity measured by NanoBRET in cells expressing WT ACKR3 or C-tail mutants (ST/A, K/A, LI/AA) and C-tail deletion mutant (ΔC). Results are presented as mean ± S.E.M of at least six independent experiments ($n \geq 6$). pEC50 values are shown; ns, not significant (one-way ANOVA with Dunnett's multiple comparison test).

**Supplementary Table 1**: Chemokines used at 100 nM to determine ligand-induced release of the HiBiT-tagged chemokine receptors shown in Figure 2.

| | | | | | | | | | |
|---|---|---|---|---|---|---|---|---|---|
| **CCR1** | CCL4 | **CCR5** | CCL3 | **CCR10** | CCL27 | **CXCR4** | CXCL12 | **ACKR1** | CCL13 |
| **CCR2-A** | CCL13 | **CCR6** | CCL20 | **CXCR1** | CXCL6 | **CXCR5** | CXCL13 | **ACKR2** | CCL5 |
| **CCR2-B** | CCL2 | **CCR7** | CCL19 | **CXCR2** | CXCL5 | **CXCR6** | CXCL16 | **ACKR3** | CXCL12 |
| **CCR3** | CCL13 | **CCR8** | CCL1 | **CXCR3-A** | CXCL10 | **XCR1** | XCL1 | **ACKR4** | CCL25 |
| **CCR4** | CCL3 | **CCR9** | CCL25 | **CXCR3-B** | CXCL10 | **CX3CR1** | CX3CL1 | **ACKR5** | CXCL10 |

**Supplementary Table 2**: Overview of antibodies used in the study.

| Name | Clone | Supplier | Catalog # | RRID | Application |
|---|---|---|---|---|---|
| **Anti-ACKR3/CXCR7** | 11G8 | R&D Systems | MAB42273 | RRID:AB_2089784 | Dot blot |
| **Anti-ACKR3/CXCR7 (APC)** | 8F11-M16 | BioLegend | 331114 | RRID:AB_10719955 | FC |
| **Anti-ACKR3/CXCR7** | 8F11-M16 | BioLegend | 331102 | RRID:AB_2089777 | FC ; dSTORM |
| **Anti-CD63 (PE)** | H5C6 | BioLegend | 353004 | RRID:AB_10897809 | FC |
| **Anti-CD63 (Pacific blue)** | H5C6 | BioLegend | 353011 | RRID:AB_10915273 | FC |
| **Anti-CD81 (PE)** | 5A6 | BioLegend | 349506 | RRID:AB_10645519 | FC |
| **Anti-CD9 (PE)** | HI9a | BioLegend | 312105 | RRID:AB_2075893 | FC |
| **Anti-CXCR4 (PE)** | QA18A64 | BioLegend | 304503 | RRID:AB_2860794 | FC |
| **Mouse IgG1 κ isotype control** | MG1-45 | BioLegend | 401402 | RRID:AB_2801451 | FC |
| **Anti-CD63 clone** | H5C6 | BD Biosciences | 556019 | RRID:AB_396297 | WB ; Dot blot |
| **Anti-CD81 clone** | 1.3.3.22 | Santa Cruz Biotechnology | sc-7637 | RRID:AB_627190 | WB |
| **Anti-calnexin** | | Cell Signaling Technology | 2433 | RRID:AB_2243887 | WB |
| **Anti-PHB1 (Prohibitin)** | 4D3G5 | Proteintech | 60092-1-Ig | RRID:AB_2252403 | WB |
| **Anti-HA.11** | 16B12 | BioLegend | 901501 | RRID:AB_2565006 | FC |
| **Anti-HiBiT** | | Promega | N7200 | RRID:AB_3665694 | WB ; Dot blot |
| **APC F(ab')$_2$ anti-mouse IgG+IgM** | | Jackson ImmunoResearch | 115-136-068 | RRID:AB_2338647 | WB ; Dot blot |
| **Anti-mouse IgG HRP (Fc-specific)** | | Jackson ImmunoResearch | 115-035-008 | RRID:AB_2313585 | WB ; Dot blot |
| **Anti-rabbit IgG HRP (H+L)** | | Jackson ImmunoResearch | 111-035-003 | RRID:AB_2313567 | WB ; Dot blot |
